# Multi-step implementation of meiotic crossover patterning

**DOI:** 10.1101/2025.11.12.687980

**Authors:** Ivana Čavka, Alexander Woglar, Yu-Le Wu, Emine Berna Durmus, Lauren Sloat, Aafke Gros, Cristina Piñeiro López, Felix Hecht, Anne M. Villeneuve, Jonas Ries, Simone Köhler

## Abstract

Crossover formation during meiosis is a tightly controlled process in which genetic information is exchanged between homologous chromosomes to increase the diversity of the progeny. In this process, an excess of double-strand breaks is introduced, but only a limited subset is ultimately processed into crossovers. Imbalances in the distribution of crossovers can lead to errors in chromosome segregation, with devastating consequences on the health of the progeny. However, the selection of which breaks are designated to become crossovers is still poorly understood, as both its timing and the ultimate molecular mechanisms are under debate. Here, we used 3D dual-color single-molecule localization microscopy and real-time confocal imaging, combined with advanced image analysis, to investigate the timing and mechanism of crossover designation in *C. elegans*. We show that meiotic crossover patterning is not established by a single decision point but depends on a dynamic, multi-layered regulation process. An initial, early selection process restricts potential crossovers to a small subset of double-strand break sites that already exhibit basic patterning features, including assurance and interference. A second, later step fine-tunes this pattern to ultimately ensure genome integrity and promote accurate chromosome segregation. Real-time imaging reveals that although the full process takes more than seven hours, key molecular events occur within minutes, highlighting how rapid local dynamic changes can give rise to an overall slow but extremely robust crossover regulation program.

## Introduction

Meiosis is a specialized form of cell division that ensures that the number of chromosomes in gametes is reduced by half, which is essential for sexual reproduction in most eukaryotes. To achieve this reduction, homologous chromosomes must first pair to align properly on the meiotic spindle, a process that critically depends on the formation of crossovers (COs), which are reciprocal exchanges of genetic material between the homologs. These COs provide the physical links necessary for accurate chromosome segregation and generate new combinations of alleles, thereby increasing the diversity across generations.

The number and placement of COs are tightly controlled: each chromosome pair must receive at least one CO to ensure proper disjunction, a principle known as CO assurance, while additional COs must be widely spaced, a phenomenon known as CO interference. Failures in this regulation can result in aneuploidy, with potentially severe consequences for fertility and viability of the offspring [1, 2].

The number, positions, and spacing of CO events are highly regulated during meiosis across sexually reproducing organisms. However, the timing of this regulation and the underlying mechanisms are poorly understood and hotly debated. Several partially contradictory models have been proposed to explain how COs are limited to a specific number. The non-random spacing of COs along chromosomes necessitates a signaling mechanism that transmits information over long distances along individual chromosomes. Competing models propose that this signal may be conveyed either through mechanical stress and force redistribution along chromosomes [3] or via the diffusion and action of molecular cues [4, 5]. The signal may act as a global inhibitor emanating from the first designated CO or through local accumulation of positive CO-promoting determinants. The models also differ in when the CO fate is determined, and in the involvement of the synaptonemal complex (SC). The SC, a 100 nm-wide protein structure bridging the homologous chromosome axes, provides both a structural scaffold for homologous alignment and a potential signaling medium. Recombination intermediates or “nodules” are consistently observed in close association with the SC, aligned above or below its midline [6]. Studies in budding yeast indicate that CO decisions occur early and independently of SC formation [7, 8], whereas studies in *C. elegans* [9, 10, 11] and various plants [12, 13, 14] imply an essential role for the SC in CO patterning. Additionally, CO patterning may also arise through multiple layered steps rather than a singular mechanism [15, 16, 17].

There are reliable cytological markers for visualizing the “end product” of CO patterning at the late pachytene stage of meiotic prophase in multiple experimental systems. These markers include for example MLH1/MLH3 foci in mammals [18, 19] and *Arabidopsis thaliana* [20], Zip3 or Hei10 foci in yeast [21] and *A. thaliana* [22], or COSA-1 foci in *C. elegans* [17]. However, testing predictions from models for CO patterning remains diffcult because we currently lack methods to visualize early signs of a CO fate among the recombination intermediates. Although many early recombination nodules are initially marked by low levels of CO markers, it is unclear which of these intermediates will mature into COs. The CO fate can be definitively identified only later, once they have accumulated high levels of CO-site markers. Here, we pursue a deeper understanding of CO patterning and CO designation using *C. elegans* oocyte meiosis.

## Results

In *C. elegans*, COs are tightly regulated and each chromosome typically receives a single CO located on one of its arms that is ultimately marked cytologically by a bright COSA-1 focus. Additionally, meiotic progression is easily tracked in this model organism because individual meiotic nuclei move sequentially through the germline, allowing us to visualize distinct meiotic stages within a single snapshot (Fig. 1A). By imaging meiotic progression in real time (see below), we measured nuclear movement at 1.1 *±* 0.35 h per row (Fig. S1), consistent with estimates from fixed samples using S-phase labeling to infer meiotic timing in *C. elegans* [23, 24]. Using this model organism, we developed two different approaches to examine when distinct properties indicative of CO patterning emerge among the cohorts of recombination intermediates present in germ cell nuclei progressing through meiotic prophase.

**Figure 1.**
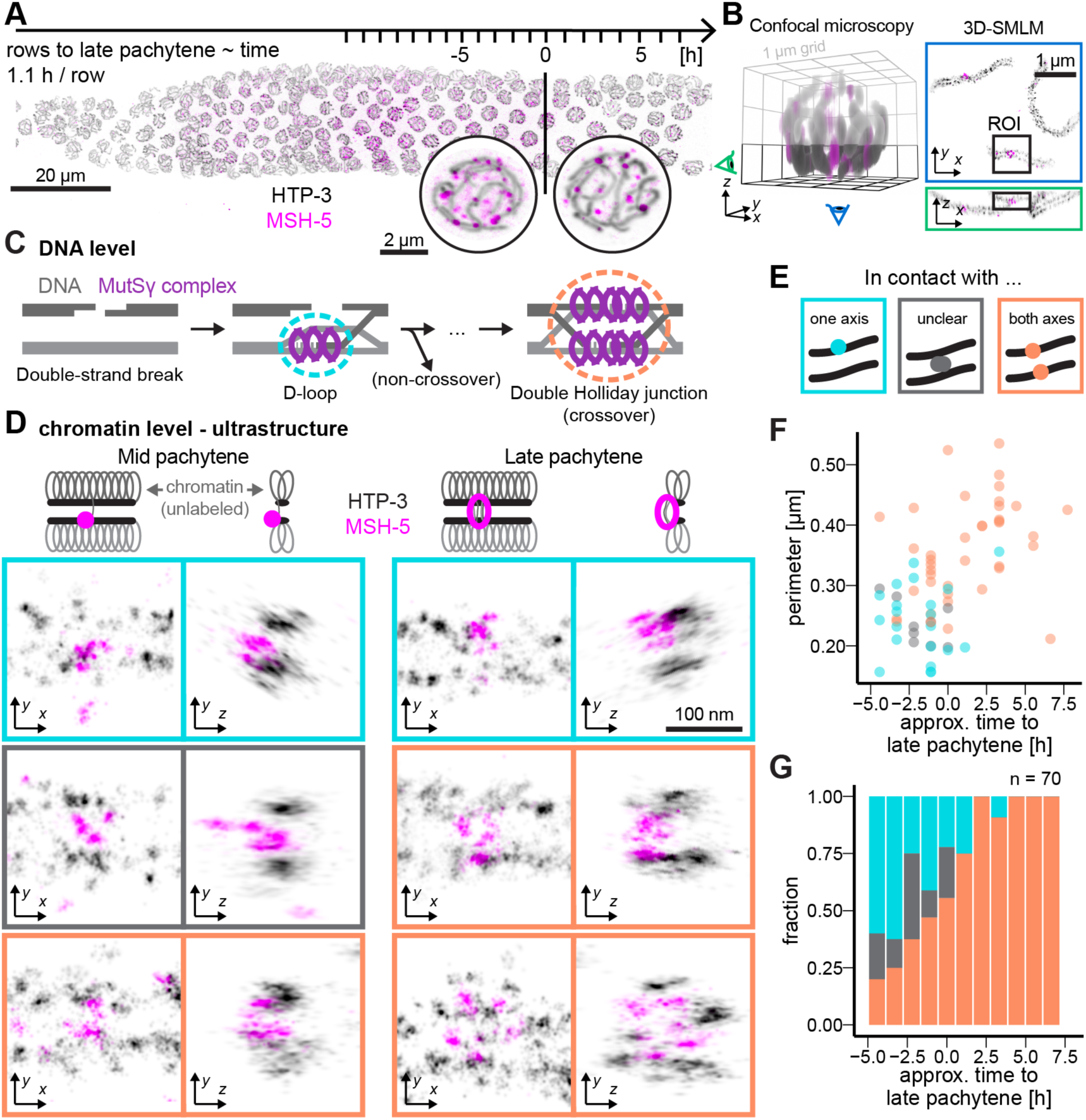
3D-SMLM reveals CO-like features in early MSH-5 recombination nodules. (A) Maximum intensity projection of pachytene meiotic nuclei in *C. elegans*. The position of nuclei within the germline is used to infer the meiotic stage of recombination nodules imaged by 3D-SMLM (single molecule localization microscopy) with nuclei progressing about one row per 1.1 hours. (B) A 3D confocal render of an entire nucleus (left) highlights the bottom imaging plane (black box), which is resolved at *↑* 30 nm resolution using 3D single-molecule localization microscopy (SMLM; right). The green and blue ”eye” icons indicate the respective viewing angles shown in the SMLM images. (C) Illustration of double strand break repair during meiosis. Early D-loop intermediates with single-end invasions may mature into non-COs or COs, while fully engaged double Holliday junctions are typically associated with CO outcomes. (D) Representative 3D-SMLM images of MSH-5 foci during mid-and late pachytene. Displayed regions of interest (ROI) are color-coded based on their contact with chromosome axes: cyan (contact with one axis), orange (contact with both axes), and gray (unclear). Chromosome axes (HTP-3) are shown in gray; MSH-5 signal is shown in pink. (E) Cartoon showing the classification of axis-contact states for MSH-5 foci. (F) Quantification of the MSH-5 focus size as measured by the perimeter of an ellipse fitted to each focus (see Supplemental Fig. S2). Color code as in (C-E). (G) Proportion of MSH-5 foci contacting one or both axes across meiotic progression. A total of 70 foci were analyzed.

### Crossover-like recombination structures form prior to the onset of late pachytene

First, by using 3D dual-color Single Molecule Localization Microscopy (SMLM) [25] to visualize the essential CO factor MutS*γ* (MSH-5) [26, 27], we reveal morphological differentiation of a subset of recombination intermediates prior to transition to the late pachytene stage (Fig. 1). MSH-5, together with its heterodimeric partner MSH-4 [28], can associate with both double Holliday junctions, which are thought to be CO-specific, and D-loop intermediates, which represent earlier intermediates that may be resolved in either COs or non-COs, both in vitro [29] and in vivo [30, 31].

From SMLM images of MSH-5 and the chromosome axis component HTP-3 in the vicinity of recombination sites (Fig. 1B), we extracted information about the ultrastructure of recombination nodules, including relative size/shape of the composite MSH-5 site and the status of association with chromosome axes (Fig. 1C-F, Fig. S2). In late pachytene nuclei, signals corresponding to MSH-5 molecules at CO-designated sites could be modeled as large ring-like structures that span both axes of the homologous chromosome pair [32] (Fig. S2, Fig. 1C-F; orange, and video S1). This pattern is consistent with previous findings of MSH-5 “doublets” at CO-designated sites in chromosome spreads [31]. These structures likely correspond to MSH-5 associated with CO-specific intermediates such as double Holliday junctions. In these structures, both ends of the double strand break engage with the homolog, which should place MSH-5 near both chromosome axes (Fig. 1C). At earlier stages, prior to the transition to late pachytene (“mid-pachytene”), recombination nodules exhibited more heterogeneity (Fig. 1D, video S2). Surprisingly, many MSH-5 structures already contacted both axes and resembled CO-designated MSH-5 sites from late pachytene. These sites likely correspond to CO-committed intermediates, and are already present in nuclei five rows or 5.5 h prior to late pachytene entry (Fig. S1). This detection of ultrastructural features characteristic of late pachytene CO-designated sites suggests that CO-biased intermediates are already present at a subset of recombination sites several hours prior to the transition to late pachytene.

By contrast, about 40% of foci in mid-pachytene were compact and associated with only one chromosome axis (Fig. 1D-F). This pattern is consistent with early-stage recombination intermediates or non-CO-destined intermediates, where only one end of the double strand break has initiated strand invasion (Fig. 1C).

### Real-time imaging of the dynamics of crossover factors

As an alternative approach to gain insight into the CO patterning process, we developed a real-time imaging assay to visualize the dynamics of CO factors associating with meiotic sites in living animals (Fig. 2). We first visualized endogenously tagged MSH-5::Halo in conjunction with COSA-1::mNeonGreen [33] as a CO-site marker. COSA-1 initially appears in mid-pachytene as numerous dim foci marking multiple recombination intermediates and later concentrates at six bright foci corresponding to designated CO sites in late pachytene [31] (Fig. 2A-B, Fig. S3, and video S3). In general, our real-time imaging mirrored the dynamic progression observed in fixed samples stained for COSA-1 and MSH-5, where early pachytene nuclei display more than six foci that gradually resolve into a stable set of six (COSA-1) or slightly more (MSH-5) foci by late pachytene, and their intensity increases over time (Fig. S3A-D).

**Figure 2.**
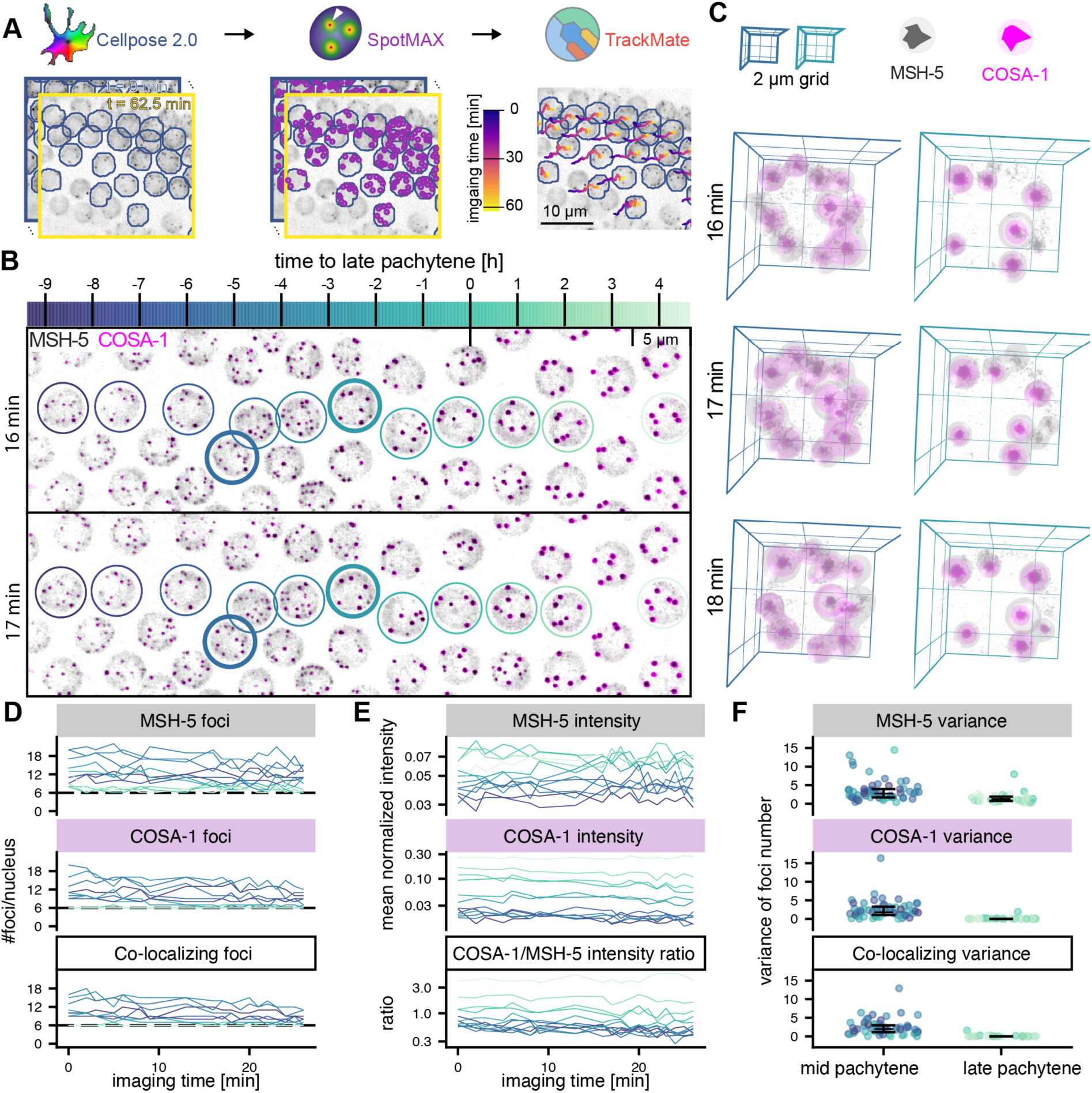
CO factors dynamically associate with recombination intermediates. (A) The automated image analysis pipeline comprises a segmentation of individual meiotic nuclei using cellpose, detection of diffraction-limited foci of crossover marker proteins using spotMAX, and tracking of individual nuclei for the duration of the acquired movie using TrackMate. (B) Maximum intensity projections of MSH-5::Halo and COSA-1::mNeonGreen at different time points show fluctuating numbers of foci in mid-pachytene nuclei, while the number of COSA-1 foci stabilizes to 6 bright, CO-designated foci in late pachytene. The time to late pachytene is calculated from the speed of the nuclei (Fig. S1). (C) 3D renders of nuclei (highlighted by thick lines in B) show dynamically appearing and disappearing foci for both MSH-5 and COSA-1 in mid-pachytene nuclei. The number (D) and intensity (E) of MSH-5 (top) and COSA-1 (bottom) foci for selected nuclei (colors as in B) are shown. (F) The variance of the number of foci is high in mid-pachytene (MSH-5: 3.6*±*2.9 (mean *±* s.d.), COSA-1: 2.5*±*2.5, co-localizing: 2.5*±*2.2; n=56 nuclei) but low in late pachytene (MSH-5: 1.6*±*1.5 (mean *±* s.d.), COSA-1: 0.1*±*0.4, co-localizing: 0.09*±*0.3; n=29 nuclei). Error bars show median *±* interquartile ranges.

Notably, our real-time imaging reveals that the number of COSA-1 and MSH-5 foci in individual nuclei fluctuates on the minute timescale (Fig. 2C-D, S3E, and video S4). Most COSA-1 and MSH-5 foci co-localize, although some foci are positive for only one marker, and MSH-5 foci typically outnumber those of COSA-1 (Fig. 2D and S3B,D). Despite the rapid fluctuations, the number of MSH-5 and COSA-1 foci very rarely drops below six - the expected number of crossovers in these nuclei - even hours before late pachytene and the first appearance of bright foci (Fig. S3B).

### Two populations of COSA-1 goci are detected approximately 7 hours before late pachytene onset

To better assess the dynamics of individual foci within meiotic nuclei and along individual chromosomes, we next imaged Halo::COSA-1 in a strain expressing endogenously tagged SYP-1::mNeongreen to visualize recombination sites in relationship to the synaptonemal complex, which assembles at the interface between aligned pairs of homologous chromosomes (Fig. 3A and video S5). The nucleus-wide dynamics of Halo::COSA-1 foci resembles the dynamics of COSA-1::mNeonGreen, suggesting that the tag does not alter the dynamic behavior of COSA-1 (Fig. S3E,F, and Fig. S4). To assess the dynamics of individual foci, we traced individual chromosomes via their synaptonemal complexes and mapped the position of recombination intermediates (Fig. 3A,B, S5A-B, and video S6). We identified two populations of COSA-1 foci in mid-pachytene nuclei. Many foci are highly dynamic, disappearing and reappearing (Fig. 3B and S5B-C, blinking foci), suggesting that COSA-1 transiently associates with these recombination intermediates. In contrast, a subset of foci remains persistently marked by COSA-1 (Fig. 3B, persistent foci). The distinction between these two populations of COSA-1 foci is evident as early as mid-pachytene and coincides with the initial appearance of COSA-1 foci (Fig. 3C). While the number of persistent foci remains constant throughout mid-pachytene, new (blinking) COSA-1 foci appear until approximately 4 hours before late pachytene (Fig. 3C, gray shaded area).

**Figure 3.**
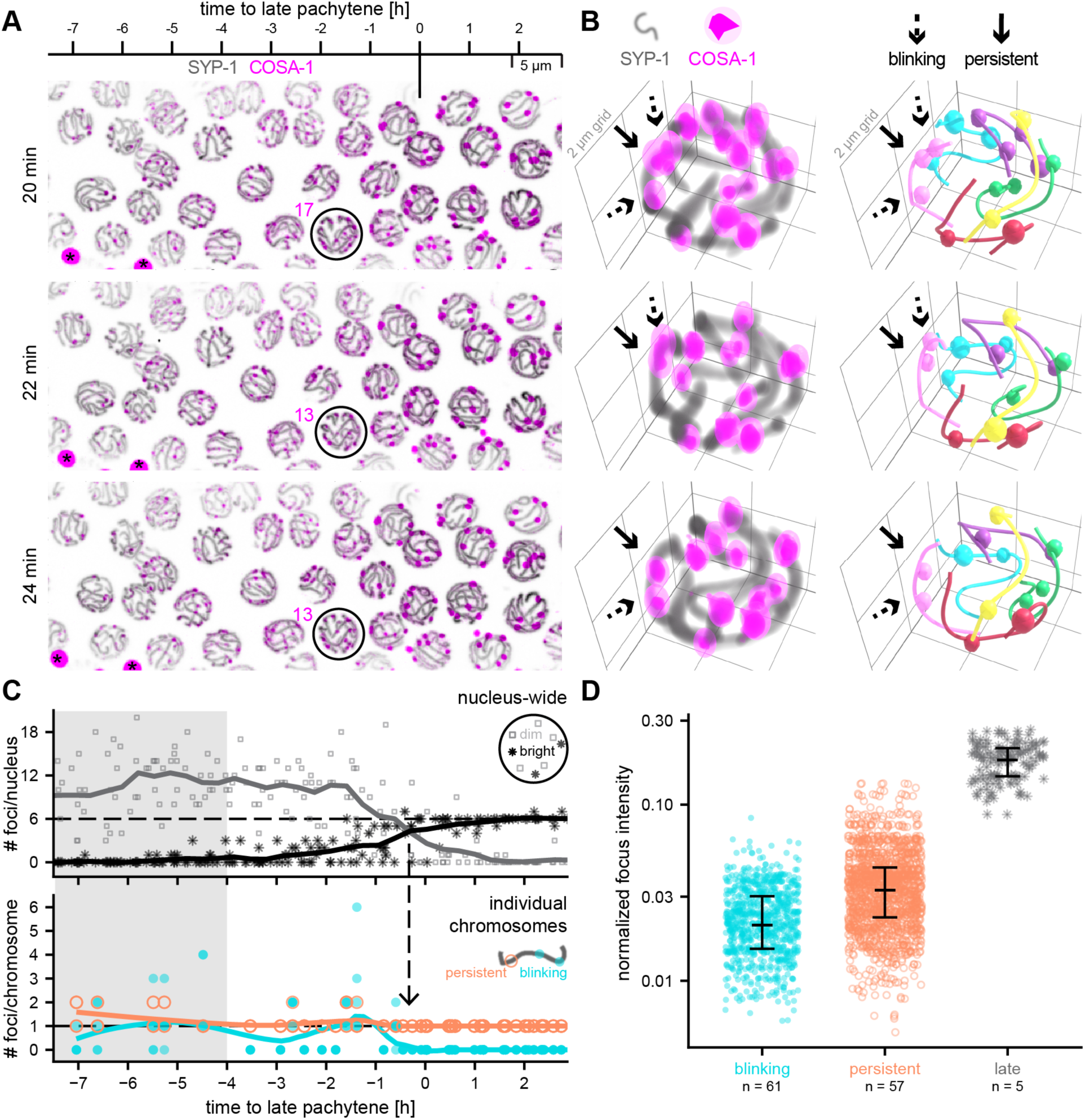
A subset of COSA-1 foci is persistent. (A) Maximum intensity projections show that the number of dim Halo::COSA-1 foci fluctuates in mid-pachytene (-7 to 0 h to late pachytene), and stabilizes to 6 bright foci in late pachytene. SYP-1::mNeonGreen stretches show the location of individual chromosomes. Black asterisks indicate highly fluorescent granules in the gut of the animal. (B) 3D renders of individual nuclei (left) show two populations of COSA-1 foci: persistent foci (solid arrows), which remain marked throughout the imaging period, and blinking foci (dashed arrows), which repeatedly disappear and reappear within minutes. Chromosome traces (colored lines) and associated spots (colored spheres) are shown on the right. The analysis was restricted to 40 frames (1 min per frame), a period over which no bleaching effects were detected (see Methods) (Fig. S8). (C) Top: Quantification of foci within individual nuclei show that variable numbers of mostly dim foci (gray squares) in mid-pachytene consolidate to mostly six bright COSA-1 foci (black asterisks) in late pachytene (n=280 nuclei from 6 different animals). Bottom: Tracing individual chromosomes revealed that each chromosome has either one or two persistent foci during mid-pachytene, beginning approximately 7 hours prior to late pachytene (43 mid-pachytene chromosomes and 5 late pachytene chromosomes were traced over time; and an additional 193 late pachytene chromosomes were analyzed at a single time point). The number of persistent foci per chromosome decreases to one shortly before late pachytene onset (arrow). The number of blinking foci is variable: blinking foci appear during early stages of mid-pachytene (gray shaded area) and decline to 0 prior to late pachytene onset. (D) While average intensities (black lines) differ between foci classes (Wilcoxon test, Benjamini-Hochberg corrected *p*-values *<* 10*^↑^*^16^ for all comparisons), there is a substantial overlap in the distributions of blinking and persistent foci. Points show intensities of spots for individual time points. n indicates the number of spots.

The number of persistent foci is relatively stable, at 1 or 2 foci per chromosome pair, during approximately 7 hours in the mid-pachytene stage, then rapidly declines to exactly one persistent focus per chromosome just prior to late pachytene. This reduction is immediately followed by the disappearance of blinking foci and a nucleus-wide increase in the brightness of all remaining COSA-1 foci (Fig. 3C, arrow).

Although the average COSA-1 intensity per nucleus increases over time (Fig. S4C), the intensity of individual foci remains variable (Fig. S5D-E). Most nuclei exhibit only dim foci during mid-pachytene and six bright foci by late pachytene. However, some nuclei show bright foci earlier than expected, or retain dim foci even in late pachytene (Fig. 3C). This heterogeneity is also reflected in the variation of foci intensities within single nuclei (Fig. S5D, lines). Similarly, the brightness of individual foci is not a reliable indicator of their persistence status. While persistent foci are, on average, brighter than blinking foci, the intensity distributions of the two populations overlap substantially (Fig. 3D), indicating that brightness alone cannot distinguish between these classes.

The presence of two populations of COSA-1 foci raises the possibility that recombination intermediates marked by blinking or persistent COSA-1 foci may represent sites destined for distinct repair outcomes. Specifically, we hypothesized that blinking COSA-1 foci correspond to intermediates destined for non-crossover repair, whereas persistent COSA-1 foci may correspond to intermediates that have acquired the potential to become crossovers. A key prediction of this hypothesis is that blinking and persistent foci will differ in their spacing and/or distribution along chromosomes, with persistent foci more closely paralleling the known behavior of COs. Thus, we set out to evaluate how the positions and spacing of these foci compared with the distributions of early meiotic recombination intermediates and of late prophase CO-designated sites.

### Crossover precursors marked by persistent COSA-1 foci partially recapitulate features of mature crossovers

Although it is well known that COs in *C. elegans* exhibit a highly biased distribution, being strongly enriched on chromosome “arms” and depleted from the centers [34], the distribution of the full cohort of earlier meiotic recombination intermediates has not been clearly established. We addressed this issue and created baselines for further comparisons by assessing the distributions of early MSH-5 foci (in mid-pachytene nuclei) and late pachytene COSA-1 foci along chromosome axes in immunofluorescence images of spread meiotic nuclei (Fig. 4A and S6A-B). The late pachytene CO foci exhibited the expected biased distribution, with 90% of foci located within 33% of chromosome length from the chromosome ends (Fig. 4B).

**Figure 4.**
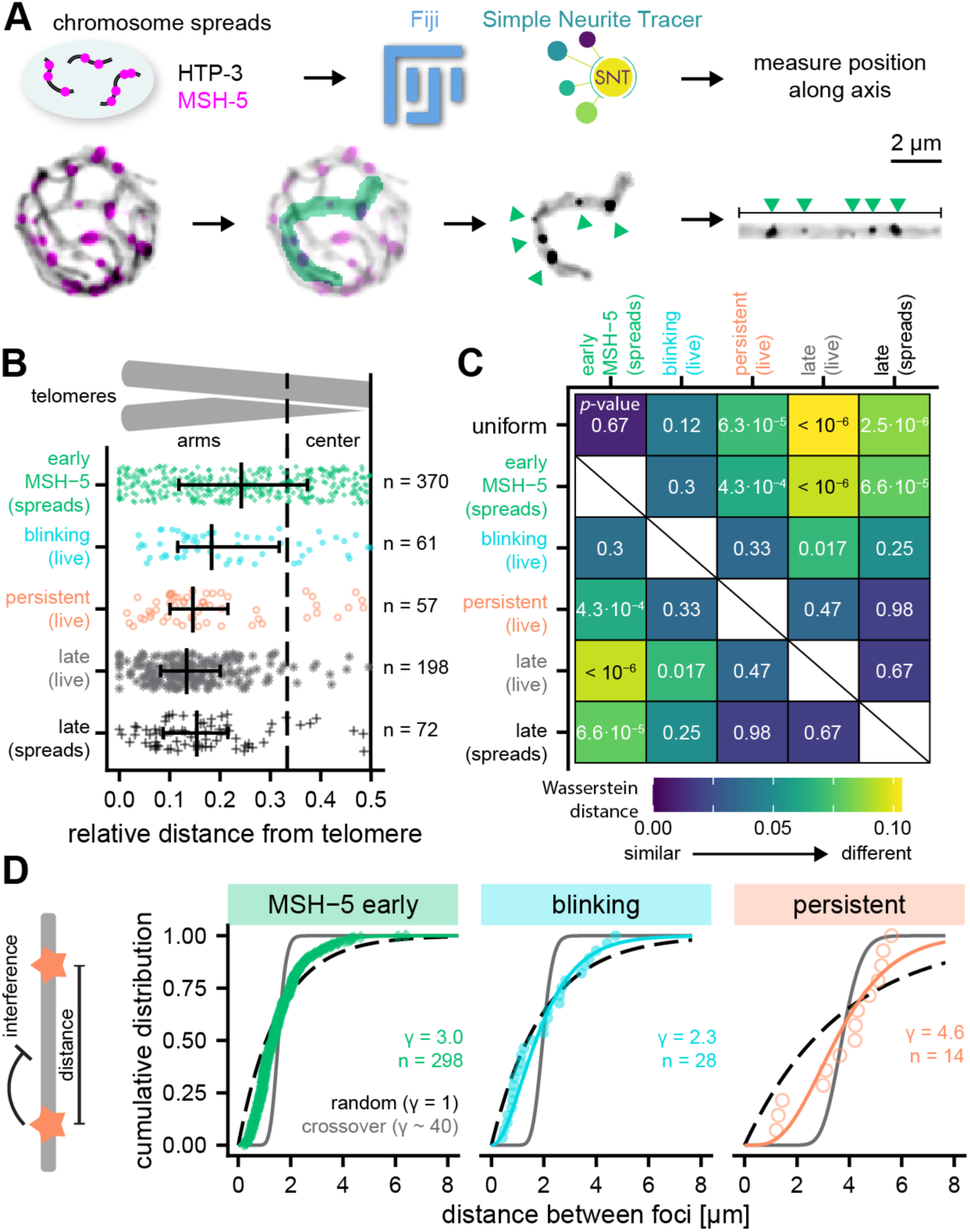
Persistent foci are CO precursors. (A) Immunofluorescence images of recombination foci and chromosome axes (HTP-3) in nuclear spreads were processed, masked, and traced in 3D using Simple Neurite Tracer to determine positions of foci along axes. (B) Distributions of foci along chromosomes, showing that persistent COSA-1 and late pachytene COSA-1 foci, observed in live imaging and spreads, are enriched on chromosome arms, while blinking foci and early MSH-5 foci in spreads show more random distributions. Data for early MSH-5 and late COSA-1 foci from spreads are shifted by 0.062 to correct for methodological differences (Fig. S6). n = number of analyzed foci. (C) Heat map of Wasserstein distances comparing the different distributions. Colors represent the extent of shift between each pair of distributions, with larger values indicating greater differences. Numbers indicate p-values for different comparisons from Kolmogorov-Smirnov tests after correction using the Benjamini-Hochberg method. (D) Fitting the distances between recombination intermediates along individual chromosomes (cartoon on the left) shows that early MSH-5 and blinking foci exhibit very little interference and resemble random distributions, while persistent foci show interference. n denotes the number of distances analyzed.

In contrast, early MSH-5 foci did not avoid chromosome centers but rather exhibited an unbiased distribution. Notably, this unbiased distribution of early MSH-5 foci, which represent post-strand exchange intermediates [31], differs markedly (Fisher’s exact test: *p <* 0.0001) from the arm-biased distribution of foci observed for DNA strand-exchange protein RAD-51 [35]. This counterintuitive finding that a marker of downstream intermediates (i.e. MSH-5 foci) exhibits an unbiased distribution despite a biased distribution of a marker of earlier intermediates (i.e. RAD-51 foci) implies that there must be center vs. arm differences in the lifetimes of early DSB repair intermediates and/or the amounts of RAD-51 loaded. This further suggests that despite the unbiased distribution of early MSH-5 foci in mid-pachytene nuclei, there may already be inherent differences in the populations of intermediates present in chromosome centers vs. arms.

Assessment of the distributions of blinking, persistent, and late COSA-1 foci from our live imaging data yielded several interesting findings (Fig. 4B-C and S6C-D). Blinking foci and early MSH-5 foci both resemble uniform distributions, whereas persistent foci and late COSA-1 foci, observed in both live imaging and chromosome spreads, exhibit significantly non-uniform distributions. In our initial analysis of distribution of late COSA-1 foci, we observed a significant difference between late COSA-1 foci in live imaging versus nuclear spreads, characterized by a systematic shift in median positions likely reflecting methodological differences in imaging and quantification approaches (Fig. S6A-F). Specifically, we find that COs are depleted from chromosome ends in spreads but not in our live imaging data set. After correcting for this methodological shift, we found that the distribution of blinking foci was indistinguishable from early MSH-5 foci, while persistent foci mimic late COSA-1 foci distributions (Fig. 4B-C).

Finally, we found that persistent foci exhibit evidence of interference. The property of CO interference, in which an (incipient) CO event inhibits the formation of others nearby to yield wide spacing between CO, is particularly strong in *C. elegans*. CO interference in *C. elegans* acts over distances longer than individual chromosomes [10, 36], so each pair typically receives exactly one CO or late COSA-1 focus. To quantify non-random spacing between foci, we estimated the shape factor of a gamma distribution describing the distances between two adjacent foci [37] (Fig. 4D) and we also calculated the normalized interference length 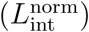 of Ernst et al., 2024 [38], which attempts to include information from chromosomes without adjacent foci (Fig. S6G). Our analysis shows that blinking COSA-1 and early MSH-5 foci display little interference, whereas persistent foci exhibit a clear interference signal using both metrics. However, the values of the interference metrics observed for persistent foci (*γ≈* 4.6, 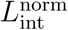 = 0.45 *±*0.13 (mean *±* s.d.) are substantially weaker than previously reported *γ* values (*γ≈* 40) [10, 11] and median 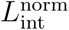 of 0.61 *±* 0.09 (Fig. S6G) for designated CO-site foci in late pachytene.

## Discussion

These findings are consistent with persistent foci representing CO-biased recombination precursors that are present at a time when the full interference pattern of CO sites has not yet been achieved. By contrast, blinking foci are destined for repair as non-COs. Therefore, a subset of DSBs or recombination intermediates is selected early for a potential CO fate in *C. elegans*. Notably, meiotic DSBs in *C. elegans* are predominantly generated during early and mid-pachytene (Fig. 5A,C). Previous studies demonstrated that DSBs formed in mid-pachytene are competent for CO formation, but not these from early pachytene [39, 40]. Thus, the timing of the formation of DSBs competent to become COs coincides with the initial appearance of COSA-1 and the establishment of persistent COSA-1 foci, and thus with the initial selection of CO-competent recombination intermediates. Indeed, a recent study found evidence that CO-competent recombination intermediates may be distinct from non-CO intermediates as early as the RAD-51-mediated strand invasion step in *C. elegans* [41]. A similar early selection mechanism concomitant with or following shortly after DSB formation has been described in other species, including fungi and mouse [42, 43, 16, 44], indicating that this regulatory step may be conserved across organisms.

**Figure 5.**
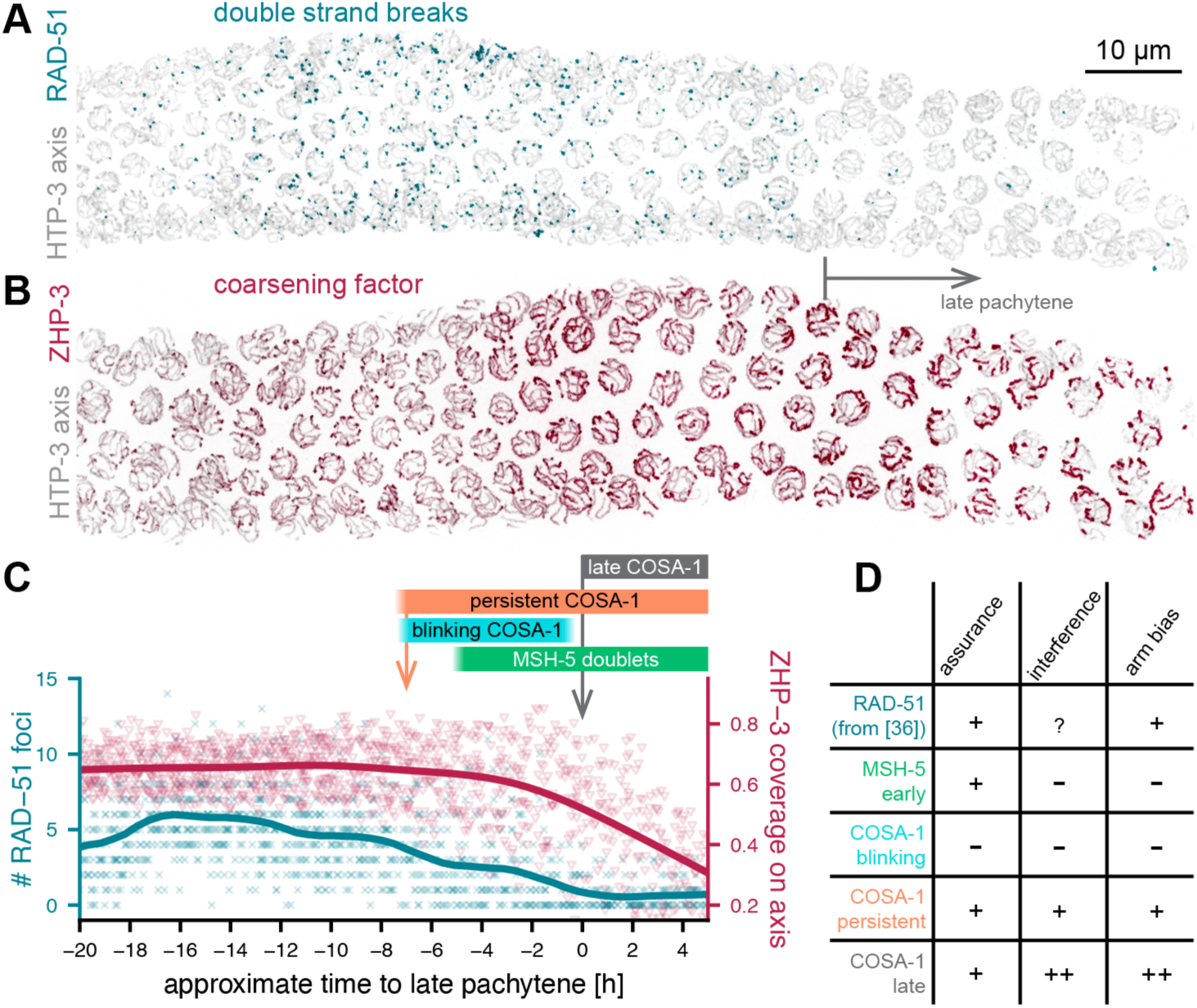
Timing of crossover designation in *C. elegans*. (A) Maximum intensity projections of V5::RAD-51 and HTP-3::HA immunostaining, showing that RAD-51 foci, a marker of early double strand break intermediates peak in number in early mid-pachytene and are removed when cells enter late pachytene (gray arrow). (B) Maximum intensity projections of ZHP-3::V5 and HTP-3::HA immunostaining, showing that ZHP-3 is initially evenly distributed along chromosomes and starts to accumulate into short patches 2-3 rows before late pachytene onset. (C) Quantification of numbers of RAD-51 foci (cyan) and coverage of the axis by ZHP-3 (red) for 8 gonads each. Colored bars above the graphs correlate the dynamics of RAD-51 and ZHP-3 with the dynamics of COSA-1 and MSH-5 foci. Both persistent COSA-1 foci as an early marker for crossover precursors (orange arrow) and MSH-5 doublets are detected before any apparent localized accumulation of ZHP-3. (D) Persistent COSA-1 foci share major hallmarks with late foci, while blinking COSA-1 foci and early MSH-5 foci do not.

The timing of this initial selection step and the dynamics of COSA-1 foci allows us to directly test predictions of models for crossover regulation. In coarsening models [4, 5], crossover precursors are selected by accumulating a CO promoting ”coarsening” factor. In *C. elegans*, this coarsening factor is proposed to be ZHP-3 [5], a RING-domain ligase that is required for CO formation [45, 46] and whose localization to the SC correlates with the ability to achieve correct crossover patterning [11]. In evaluating the distribution of ZHP-3 along the chromosome axes, we found that ZHP-3 remains uniformly loaded along the axes throughout most of mid-pachytene (Fig. 5B,C). Thus, the initial selection step, as evident by persistent COSA-1 foci in live imaging and by presence of MSH-5 doublets in 3D-SMLM images, precedes any obvious coarsening behavior of ZHP-3 and is thus inconsistent with the coarsening model. Furthermore, out data violate another prediction of coarsening models, namely that the intensities of neighboring foci should be negatively correlated, as material would redistribute from one focus to another. Since COSA-1 and ZHP-3 directly interact [47], this negative correlation is also expected for COSA-1 foci. However, we observe no negative correlation, either along individual SCs or in 3D across the nucleus, which further argues against a diffusion-driven competition mechanism (Fig. S7). Instead, the pronounced fluctuations in COSA-1 and MSH-5 intensity, at both blinking and persistent foci, may reflect dynamics in homologous partner interactions. Such instabilities could underlie the frequent template switching observed in both CO and non-CO repair outcomes in several organisms [48, 49, 50].

The existence of an early, interfering selection step raises the possibility that CO regulation involves rapid, chromosome-wide communication. Since this selection likely occurs concomitant with or shortly after DSB formation [39, 40, 41], such communication must act very quickly, and the selection of one CO-competent break must suppress the formation of others nearby. The required speed likely exceeds what can be achieved by diffusion or enzymatic signaling alone, but could instead point to a mechanical mechanism, such as stress propagation, tension-sensitive feedback, or dynamic structural transitions within the SC or chromosome axes [3, 51, 52].

The early selection step establishes crossover assurance, as it results in at least one persistent COSA-1 focus on each pair of chromosomes (Fig. 5D). However, the initial selection is imperfect, and early persistent foci exhibit weaker arm bias and weaker interference than late COs. This discrepancy indicates that a second regulatory step is necessary to achieve the precise pattern of exactly one CO per chromosome that is typically positioned on the arms. The fact that we observe chromosomes with two persistent foci until 1.5 hours before the entry into late pachytene but not later implies that this second step must occur during this time window (Fig. 3C). Notably, this time window coincides with first signs of coarsening of ZHP-3 (Fig. 5). This second regulatory step must also increase the arm bias. We therefore propose that this second step is likely to retain the more distal persistent foci. However, COSA-1 intensity does not correlate with the position of the persistent foci along the chromosome, and when two persistent foci are present the more distal focus is not consistently brighter than its more central counterpart (Fig. S7C). Thus, accumulation of material is unlikely to be the determining factor in selecting the final CO site. Instead, we propose that a length-sensing mechanism may select which persistent focus becomes the ultimate CO. In *C. elegans*, the CO position determines the large-scale reorganization of the chromosome axes and the SC [53], and this reorganization is likely implemented through a diffusion-based length-sensing mechanism [54]. Thus, length-sensitive removal of factors such as ZHP-3 may help select the final CO. Importantly, we observe a final increase in COSA-1 intensity only after this second step (Fig. S5D-E). We speculate that a checkpoint mechanism enforces this transition. This checkpoint may activate only when nearly all non-CO-destined intermediates (i.e., blinking foci and supernumerary central persistent foci) have been resolved, leaving only persistent CO-designated sites. Such a nucleus-wide transition could be mediated by inactivation of CHK-2 kinase, which was previously shown to coincide with the increase in COSA-1 intensity [55].

Rather than a single, unifying model such as coarsening, we propose that CO patterning emerges from a hierarchical and possibly modular system of regulatory layers. Whether these layers represent a single integrated physical mechanism or represent distinct processes remains to be determined. Such a multi-layered regulation of CO events may also occur in other species, where evidence suggests several consecutive steps that convert DSBs into interfering, CO-competent intermediates that ultimately mature into strongly interfering COs [16, 15]. Together, our findings support a flexible and evolutionarily conserved framework in which CO number and position are jointly controlled through the integration of mechanical, spatial, and molecular cues.

## Materials and Methods

### *C. elegans* maintenance and CRISPR/Cas9 genome editing

*C. elegans* animals were maintained under standard conditions at 20*^↓^*C on standard nematode growth medium (NGM) agar inoculated with *E. coli* OP-50 [56]. All experiments were performed with 18-24 h post-L4 animals. *C. elegans* strains used in this study are listed in Table S1.

CRISPR/Cas9-mediated genome editing was performed by micro-injection of pre-assembled CRISPR/Cas9 complex into the gonad of adult worms following previously reported procedures [57, 58]. Reagents for individual CRISPR/Cas9-mediated genome edits are listed in Tables S2 - S3. All genome editing experiments were designed using sequence information obtained from wormbase [59]. Guide RNAs were identified with CCTop - CRISPR/Cas9 target online predictor [60, 61].

Briefly, worms were injected with a mix containing 1.5 µM of Cas9 enzyme (Alt-R^™^ S.p. Cas9 Nuclease V3, IDT, 1081058) in complex with a specific guide RNA, and varying concentrations (Table S2) of a target-specific repair template diluted in nuclease-free water (AM9939, Thermo Fisher Scientific) or duplex buffer (IDT, 11-01-03-01). Additionally, the mix contained 5 ng/µL of pCFJ90, and 2.5 ng/µL of pCFJ104 plasmids that served as co-injection markers [62]. Successful genome editing events were identified by PCR (Table S3) and confirmed by Sanger sequencing (Eurofins Genomics Germany GmbH).

To assess the fidelity of meiosis in all generated strains, their fertility was assessed by brood counts. For each strain and the wild-type isolate (Bristol N2), six hermaphrodite worms at the L4 larval stage were picked on six OP50 NGM agar plates. These P0 animals were transferred to fresh plates every 24 hours for five days. Laid eggs and hatched L1 larvae were counted immediately after each transfer to determine the brood size. Once the progeny reached adulthood, we determined the number of surviving and male progeny. We determined the viability of the progeny (the total count of surviving progeny divided by the total brood size) and the incidence of males (the total count of male progeny divided by the total count of surviving progeny) for each initial P0 animal.

A comparison of the mean viabilities of the progeny and incidence of males of each assessed strain and the SMN260 strain (*cosa-1* (*ske25* [*HaloTag::cosa-1*]) III) was done using a two-sided Wilcoxon rank sum test using the *compare means* function from the *ggpubr* (version 0.6.0) package in R (version 4.4.3).

### *In vivo* imaging of the *C. elegans* germline

**Sample preparation.** We used *C. elegans* strains expressing either SYP-1 C-terminally tagged with an mNeonGreen [63] fluorescent protein and the pro-crossover factor COSA-1 N-terminally tagged with HaloTag7 [64], or a strain expressing COSA-1 C-terminally tagged with mNeonGreen and MutS*γ* complex component MSH-5 C-terminally tagged by HaloTag9 [65]. Additionally, to compromise the locomotion in response to blue light, the *gur-3* gene deletion (*ok2245*), and a point mutation in *lite-1* (*ce314*) were introduced through crosses [66].

To visualize the HaloTag-labeled endogenous proteins, we grew worms on bacterial lawns that contained HaloTag ligand. To this end, we pelleted 1-3 mL overnight OP-50 culture, washed the pellet with M9 buffer (22 mM KH_2_PO_4_, 42 mM Na_2_HPO_4_, 85.5 mM NaCl, 1 mM MgSO_4_), and resuspending the pellet in 100 µL of M9 buffer containing 0.2% Tween^®^ 20 and 10 µM HaloTag ligand conjugated with a JF639 organic dye [67]. Bacterial lawns were prepared by inoculating individual NMG agar plates with 20-30 µL of resuspended OP50 bacteria. Additionally, to contain worms on the bacterial lawn, a palmitic acid ring was imprinted around the bacterial lawn [68]. L4 larvae were picked from plates that were well-fed for several generations. 15-22 hours before imaging, the worms were transferred to fresh NGM agar plates inoculated with HaloTag ligand.

For long-term imaging of *C. elegans* meiosis *in vivo* we used a modified mechanical immobilization method [69]. Briefly, we cast 40 µL of 7% (w/v) agarose (Invitrogen UltraPure^™^ Agarose-1000, 16550100, ThermoFisher Scientific) gel pads between a vinyl record and a glass slide spaced by tape as described [69]. The pad was incubated in an agarose soaking solution (20 mM serotonin creatinine sulfate monohydrate (Sigma-Aldrich, H7752-1G), 0.002% (w/v) of tetramisole hydrochloride in M9 buffer) within a humid chamber for 30 min to two hours. Worms were mounted within the channels of an agarose pad (approx. 3 mm x 5 mm). To prevent oxygen depletion, the pad was enclosed in a sandwich of a 12 mm coverslip and an imaging dish with a glass bottom (µ-Dish 35 mm, high Glass Bottom, Ibidi, 81158) that were held together by droplets of M9 buffer placed around the agarose pad without any additional sealing. To prevent sample desiccation, a wet tissue ring was placed around the plastic edge inside the dish, away from the agarose pad. All solutions were prepared fresh.

### Image acquisition

Real-time images of *C. elegans* germlines were acquired at 20*^↓^*C with an Evident IXplore SpinSR spinning-disk confocal microscope equipped with a UP-LSAPOS100X (NA 1.35) silicon oil immersion objective and a CSUW1-TS 2D 50 µm spinning disk. Both fluorescent reporters were excited simultaneously by a 488 nm laser at 4-7% power, and a 640 nm laser at 40-50% power. A 561 nm long-pass dichroic mirror was introduced to the optical path to split the emitted fluorescence onto two separate sCMOS cameras. Additionally, the light of the reflected and transmitted channels was passed through a 525/50, and a 685/40 band-pass filter, respectively. The microscope was controlled through the CellSens Dimension software (version 3.2, Evident).

### Segmentation and tracking of individual meiotic nuclei

To segment the meiotic nuclei in 3D over time, we used cellpose 2.0 [70, 71] with a model specifically trained for segmentation of meiotic nuclei within the *C. elegans* germline [72, 11]. The segmentation was performed on the channel containing either the images of SYP-1::mNeonGreen or MSH-5::HaloTag9 processed by a gaussian blur filter (*ε* = 3.5 px or *ε* = 1.5 px respectively) prior to segmentation.

To remove partial and joint nuclei, aberrant structures, or crescent-shaped leptotene/zygotene nuclei, the segmented nuclei were filtered based on their size and sphericity: We only considered nuclei with a volume corresponding to spheres of a diameter between 3.5 and 8 *µ*m, and sphericities greater than or equal to 0.75. Sphericities were approximated from the surface of a mesh generated from the individual 3D object using *scikit-image* (version 0.19.3) [73] in Python (version 3.9.15).

To track meiotic nuclei over time, we used the Fiji Plugin TrackMate [74, 75]. Specifically, we used a linear assignment problem (LAP) tracker after a correlation-based alignment of the movie frames to correct for animal movement. Trackmate parameters were altered depending on the movement of the imaged *C. elegans* animal. Example values for parameters are listed in Table S5. All tracking results were inspected and manually corrected as necessary.

Additionally, frames where the animal moved during the *z* -stack acquisition were identified by inspecting the movies in Fiji. All such frames are excluded from the reported results.

### Foci detection

Detection of diffraction-limited foci within each cellpose 2.0 identified object (meiotic nucleus) was done by the spotMAX software [33]. The parameters used to identify COSA-1::mNeonGreen, MSH-5::HaloTag9, and HaloTag7::COSA-1 foci are listed within the corresponding Supplemental files (S1 COSA-1-mNG.ini for COSA-1::mNeonGreen, S2 MSH-5-Halo.ini for MSH-5::HaloTag, and S3 Halo-COSA-1.ini for HaloTag7::COSA-1, respectively).

For HaloTag7::COSA-1 foci detection, the initial thresholding step of spotMAX was replaced by generating masks with a custom-trained ilastik [76] pixel classifier.

To this end, the SYP-1::mNeonGreen channel was aligned to the far-red HaloTag7::COSA-1 channel by a custom Python script based on the correlation of maximum intensity projections of individual frames of the movie for offset correction. The correction was performed consistently for all time frames in all three directions using the mode of estimated shift values. Subsequently, the ilastik classifier was trained on several selected frames and applied to the remainder of the offset-corrected frames (*z* -stacks) to distinguish between signal and background.

### Data processing

The spotMAX output was further processed by custom R (version 4.4.3) and Python (version 3.9.15) scripts. Specifically, identified foci with a small difference between the signal of a detected focus and the background (Hedge’s effect size below 1.1, 2.1, or 3.0 for HaloTag::COSA-1, MSH-5::HaloTag9, and COSA-1::mNeonGreen, respectively) were discarded. The foreground integral estimated from a 3D Gaussian fit to each identified focus was used to estimate foci intensity. Additionally, raw spotMAX foreground estimations were adjusted for photobleaching by histogram matching and those values are used synonymously with intensity in this article.

To estimate the meiotic progression, we determined the position of each nucleus relative to the inflection point of the increase in the intensity of COSA-1 foci that occurs at the transition from mid to late pachytene.

Firstly, using principal component analysis (PCA) on the centroids of the segmentation masks, the germlines were rotated such that the axis of meiotic progression is parallel to the *x* -axis. Second, the central line of the gonad was approximated by locally weighted scatterplot smoothing (*lowess* function from the *statsmodels* (version 0.13.5) Python library) of the rotated coordinates of the centroids. Lastly, the logarithm of the median normalized amplitude of the fitted gaussian peaks (A fit in spotMAX output) within each object along the centerline was used to determine the direction of meiotic progression and the point of late pachytene entry. Specifically, COSA-1 foci intensity was fitted to a cumulative gaussian function to determine the inflection point (onset of late pachytene) using a custom R script relying on the *minpack.lm* (version 1.2-4), *numDeriv* (version 2016.8-1.1), *reshape2* (version 1.4.4), and *tidyverse* (version 2.0.0) packages. The position of each nucleus along the centerline was adjusted to represents its distance to late pachytene onset (Fig. S1A,B).

Assigning the nuclei with a position relative to the onset of late pachytene allowed us to estimate the speed of nuclei movement as a slope of a linear model fitted to the relationship between the estimated position within the gonad and the imaging time. Additionally, to estimate the meiotic progression in fixed gonads, we estimated the distance between rows of nuclei by calculating the distance between the centroid coordinates of the nearest neighboring nuclei (Fig. S1C-E).

To prevent the confounding effect of photobleaching on the conclusions about foci dynamics, we only assessed the dynamics of recombination factors within the first 40 frames of the movie (Fig. S8). The cutoff frame was determined by comparing the distribution of the median foci number along the gonad to that in the first frame using a Wilcoxon signed-rank test corrected by the Benjamini-Hochberg procedure.

To assess the difference in the median foci number or variance in foci number per nucleus between HaloTag::COSA-1 and COSA-1::mNeonGreen, we used generalized additive mixed models using the R package *mgcv* (version 1.9-3) by applying a smoothing function to the number or variation of foci within individual nuclei during pachytene. Two models were fitted: one allowing the smooth term to vary by genotype (HaloTag::COSA-1 vs. COSA-1::mNeonGreen), and one assuming a shared smooth term across genotypes. The difference between models was assessed using an ANOVA test on the fitted models to test for a significant genotype-specific effect.

### Synaptonemal complex tracing and foci tracking

Synaptonemal complexes of nine mid-pachytene and a single late pachytene nuclei were manually traced using *BigTrace v.0.2.1* plugin for Fiji (https://github.com/ekatrukha/BigTrace).

The position of a COSA-1 focus along the synaptonemal complex was approximated using a custom Python script:

Raw traces from BigTrace were smoothed using the spline interpolation from *SciPy* (version 1.9.2). Equidistant points (120 nm spacing) were sampled along the processed traces using a Python function based on the MATLAB function *interparc* (https://www.mathworks.com/ `matlabcentral/fileexchange/34874-interparc`). Next, the offset-corrected foci coordinates (see section *Foci detection*) were assigned to the nearest synaptonemal complex trace based on Euclidean distance.

Importantly, when tracing the synaptonemal complex over time, the start of the trace was kept consistent at a single end of the synaptonemal complex. This allowed us to identify the regions of the synaptonemal complex with a persistent or transient association of the COSA-1 signal.

To determine the position of a focus along the trace, we measured the cumulative segment length from the start of the trace to the point associated with the focus. In each time frame, an interval was defined around this position using a width determined by the resolution limit and the orientation of the synaptonemal complex relative to the imaging axis. Overlapping intervals across multiple frames were grouped and tagged as containing *persistent* or *blinking* foci. The classification of COSA-1 focus-associated regions along synaptonemal complex traces was performed with a custom R script using *GenomicRanges* (version 1.58.0), *IRanges* (version 2.40.1), and *mclust* (version 6.1.1).

Additionally, we disregarded the synaptonemal complexes (12 out of 60) in which individual foci could not be reliably tracked due to the synaptonemal complex’s orientation relative to the imaging direction. Due to lower resolution along the *z* -axis, even upon manual inspection of these chromosomes, we were unable to reliably determine the behavior of foci within certain regions aligned with the *z* -axis. Notably, as it is significantly shorter than autosomes preventing an accurate assignment of foci, the X chromosome was always excluded from the analysis.

### Arm bias and interference evaluation

To evaluate the arm bias, we compared the distances of foci to its closest end of the trace as a proxy for distal (close to telomere) or central location. Positions on opposite arms were mirrored around the chromosome midpoint so that all values ranged from 0 (telomere) to 0.5 (center). To correct for systematic offsets between live imaging and spread in situ experiments, we rescaled the focus positions based on the median difference between late COSA-1 foci in live versus spread samples (Fig. S6). Distributions were compared using the Wasserstein distance, which is a measure of the distance required to align one distribution with another and assess shifts between them. p-values were calculated using a Kolmogorov-Smirnov test. Interference between foci was analyzed by comparing the normalized interference length as described in [38] using a Mann-Whitney test. Additionally, we estimated the gamma shape by fitting a gamma distribution to the distances between two or more foci on the same chromosome using the *fitdistrplus* R package (version 1.2-2). Furthermore, to investigate the relationship between the intensities of neighboring foci, we paired each COSA-1 focus with its closest neighbor along the synaptonemal complex trace (1D distance), or within the nucleus (Euclidean 3D distance). For each pair, we calculated the Pearson correlation coe”cient using R. For blinking foci, we set the intensity value to zero in the frames when they were not detected.

### Image and data visualization

Images showing individual movie frames are maximum intensity projections of the originally acquired *z* -stacks smoothed by Gaussian blur (*σ*= 1 px) prepared in Fiji [77]. 3D-rendering was done using Blender and the Microscopy Nodes add-on [78]. Microcopy data was rendered as volumes, and segmented spots and nuclei are visualized as surface renders. All data visualization was done using functions from the *ggplot2* (version 3.5.1) package in R.

### Imaging of the chromosome spreads

#### Sample preparation

Partial spreading of *C. elegans* gonads was performed as in Pattabiraman et al., 2017 [79] and Woglar et al., 2018 [31] with slight modifications. Specifically, 500 bleach-synchronized worms (24-36 hours post L4) were dissected on an ethanol-washed 22 × 40 mm coverslip (No. 1.5, VWR, 48393 172) in 10 µL dissection solution (water supplied with 0.1% Tween-20). 50 µL spreading solution (32 µL of fixative (4% w/v Paraformaldehyde and 3.2–3.6% w/v Sucrose in water), 16 *µ*L of Lipsol solution (1% v/v/ Lipsol in water), 2 *µ*L of Sarcosyl solution (1% w/v of Sarcosyl (Sigma-Aldrich, L-5125) in water) were slowly dropped onto the dissection mix (worm debris and non-cut worms were not removed) before distributing them over the whole coverslip with a pipette tip. Coverslips were left to dry at room temperature for 2 hours to overnight before being washed for 20 minutes in methanol at -20 *^↓^*C, and then rehydrated by washing three times for 5 minutes in PBST (0.1% Tween-20 in PBS). A 20-minute blocking in 3% w/v BSA in PBST at room temperature was followed by overnight incubation with primary antibodies at 4°C (antibodies diluted in 3% w/v BSA in PBST supplied with 0.05% w/v NaN3). Details on the *C. elegans* strains and antibodies are listed in Table S6.

Finally, coverslips were washed three times for 5 minutes in PBST before incubation with secondary antibody for 2 hours at room temperature. After PBST washes, the nuclei were immersed in Vectashield (Vector Laboratories, H-1000-10), and the coverslip was mounted on a slide and sealed with nail polish. Imaging was performed using the DeltaVison OMX Blaze microscopy system as previously described [31]. Images of MSH-5 and HTP-3 stained mid-pachytene nuclei were acquired with widelfield illumination, images of COSA-1 and HTP-3 in late pachytene with structured illumination microscopy (SIM).

#### Assessing the distribution of MSH-5 foci along the synaptonemal complex

The widefield images of nuclei stained for MSH-5 and HTP-3 (deconvolved and corrected for registration using SoftWoRx) or SIM images of nuclei stained for COSA-1 and HTP-3 (3D-reconstructed and corrected for registration using SoftWoRx) were transformed first into RGB color images (with the brightness and contrast of the foci signal being enhanced) and then into monochrome 8-bit images, in which MSH-5/COSA-1 foci appeared as bright dots on dimmer HTP-3 lines. In such images, individual chromosomes were traced using the Simple Neurite Tracer plugin of ImageJ [80]. First, chromosomes were traced end-to-end, followed by tracing each MSH-5/COSA-1 focus from one end.

### Single-molecule localization microscopy

#### Sample preparation

We used a *C. elegans* worm strain expressing MSH-5 (MutS*γ* complex component) C-terminally tagged with a V5 epitope tag, HTP-3 (chromosome axis component) C-terminally tagged with a hemagglutinin (HA) epitope tag, and COSA-1 N-terminally tagged with HaloTag7. Age-matched *C. elegans* animals fed with JaneliaFluor 503 conjugated HaloTag ligand were dissected, fixed, and stained as previously described [81].

To label MutS*γ* complex and chromosome axes, we used 1:200 dilution of anti-V5 (Thermo Fisher Scientific, R960-25, mouse monoclonal) and anti-HA (Thermo Fisher Scientific, A190-138A, goat polyclonal). Incubation with primary antibodies was done overnight at 4*^↓^*C, and incubation with secondary antibodies was 1 to 2 hours at room temperature using a 1:50 dilution of F(ab’)2-anti-mouse (Jackson Immunoresearch, AB 2340761, donkey polyclonal) conjugated with AlexaFluor 647 (ThermoFischer Scientific, A37573) and F(ab’)2-anti-goat (Jackson Immunoresearch, AB 2340386, donkey polyclonal) conjugated with CF660C (Biotium, 92137) or CF680 (Biotium, 92139) organic dye. All antibody dilutions were prepared in 1XRoche blocking buffer (Roche, 1096176001). The samples were imaged within a week of immunolabeling.

#### SMLM acquisition, image reconstruction, and post-processing

All SMLM acqui-sitions were performed at room temperature using a custom-built 3D-SMLM microscope [82] equipped with UPLSAPO100XS (NA 1.35) silicon oil objective and operated via *µ*Manager and htSMLM [83] as previously described [81].

SMLM image reconstruction [25] and post-processing (channel assignment and drift correction) were performed using the SMAP software (Super-resolution Microscopy Analysis Platform) [84], following previously reported procedures [25, 85, 86, 87, 88]. Furthermore, dim and out-of-focus emitters were removed by removing the localizations with *z* -coordinate outside of a range from -300 to 300 nm, or with a localization precision above 15 nm in lateral (*xy*) and 25 nm in axial (*z*) direction. The super-resolved images were rendered in the SMAP software [84]. Each localization was represented by a gaussian with a width proportional to its localization precision and the color based on the assigned channel [89, 90, 86, 85, 91]. The quality of the SMLM images was assessed by calculating the Fourier ring correlation (FRC) resolution [92, 93] using the FRC resolution plugin in SMAP.

### Identification of recombination intermediates in 3D-SMLM images

To identify recombination intermediates in 3D-SMLM data, the localizations corresponding to the AlexaFluor647 channel of MSH-5 within 350 nm around the center line of the neighboring chromosome axes (HTP-3) localizations were analyzed using the DBSCAN clustering algorithm. To perform clustering, we used the *dbscan* MATLAB function with *epsilon neighborhood* of 100 nm and *minimum number of neighbors required for a core point* set at 20. The identified clusters were additionally filtered based on the distance of their center to the centerline of the chromosome axes. Only clusters within 120 nm of the axes centerline were identified as possible recombination intermediates and further inspected. Clusters that were potentially cut when filtering out localizations outside of the *±* 300 nm range from the focus plane (out-of-focus emitters filter), or clusters corresponding to background (nucleoplasmic) signal were excluded from the analysis and reported results.

The sites of MSH-5 localizations within a 400 nm square around the cluster’s center were annotated and fitted with the elliptical model described in the following section. All identified sites were manually inspected and classified into categories depending on whether they contact one or both chromosome axes (Fig. 1). Additionally, to approximate timing to the late pachytene, we assigned the foci of each nucleus with the corresponding row to the late pachytene multiplied by the estimated time spent in each row (ratio of mean nucleus distance and mean speed of nuclei = 1.1 h/row, Fig. S1).

### Maximum likelihood fitting of the ellipsis

From the recognized recombination intermediates (section *Identification of recombination intermediates in 3D-SMLM images*), we selected individual regions of interest containing the chromosome axes in a frontal to slightly tilted view in dual-color SMLM images with MSH-5 in the AlexaFluor647, and HTP-3 in the CF660C/CF680 channel.

Subsequently, localizations in the AlexaFluor647 channel were described by maximum likelihood fitting of an elliptical model [94]. The fit was initialized by an elliptical model (Table S7) with a center in the median coordinates of MSH-5 localizations within the region of interest. Major and minor spans (a and b in Fig. S2A) of the model were optimized while allowing the model to be translated within the region of interest and rotated around all three axes of the coordinate system. Translation and rotation of the model are described by extrinsic parameters integral to the LocMoFit framework [94]. The fitted ellipses’ perimeter from regions of interest where the fit converged, and ellipses engulfed the MSH-5 localizations (Fig. S2), was used to compare the size of recombination intermediates. Statistical comparison of perimeter distribution between different categories of MSH-5 foci with the Kruskal-Wallis test, followed by Wilcoxon rank-sum test with Benjamini-Hochberg p-value correction, was performed using functions from *rstatix* (version 0.7.2).

### Confocal imaging of extruded and immunostained *C. elegans* germlines

#### Sample preparation

Samples were prepared in the same manner as for 3D-SMLM, with the exception of the use of the secondary antibodies and the mounting technique. Namely, after the overnight incubation with 1:1000 dilution of the primary antibodies, samples were incubated for 1 hour at room temperature using a 1:1000 dilution of donkey-anti-mouse polyclonal IgG conjugated with AlexaFluor 546 (Invitrogen, A10036) and donkey-anti-goat polyclonal IgG conjugated with AlexaFluor 647 (Jackson Immuno Research, AB 2340863), or a 1:1000 dilution of donkey-anti-mouse polyclonal IgG conjugated with AlexaFluor 546 and donkey-anti-goat polyclonal IgG conjugated with AlexaFluor 488 (Invitrogen, A-11055). Following the DNA staining in DAPI (Sigma-Aldrich/Merck, D9542-1MG) 0.1 µg/mL solution, the samples were mounted between a glass slide (Fisherbrand Microscope slides T/F

Ground 0.8-1.0 mm thick, Fisher scientific, 7107) and a coverslip (Circular cover glass 18 mm No. 1.5, Carl Roth, LH23.1) in 18 µL of mounting medium (Invitrogen ProLong Glass Antifade Mountant, ThermoFischer Scientific, P36982). Samples were kept overnight at room temperature, protected from light, to cure the mounting medium.

### Image acquisition

Images of entire *C. elegans* germlines were acquired as multiple consecutive, partially overlapping, *z* -stack tiles on an Evident IXplore SpinSR spinning-disk confocal microscope equipped with a PLAPON 60X (NA 1.42) oil immersion objective combined with 3.2X post-magnification, and a SoRa spinning disk. The microscope was controlled through the CellSens Dimension software (version 3.2, Evident).

### Image quantification

Maximum intensity projections of individual tiles were stitched using the Fiji ”Grid/Collection stitching” plugin [77]. Nuclei were segmented in 3D stacks using a custom cellpose 2.0 [71] model on the DAPI channel as described [11]. To linearize the gonads, the *XY* -coordinates of nuclei centers in stitched images were first rotated along the principal axis of the gonad using a linear fit. The position of each nucleus along the gonad was defined by fitting a LOWESS curve through the rotated centroids to extract their 1D positions. RAD-51 foci were counted automatically in 3D using spotMAX as described [33]. To determine the localization of ZHP-3 along the chromosome axes, 3D images of HTP-3 and ZHP-3 were analyzed in Python using *numpy*, *pandas*, and *scikit-image*. For each nucleus, HTP-3 and ZHP-3 signals were thresholded using the Otsu method to identify positive pixels. The ZHP-3 coverage on the axis was defined as the fraction of ZHP-3–positive pixels located along the HTP-3–positive chromosomal axis. Values were plotted along the linearized gonad to provide a measure for the distribution of ZHP-3 along the chromosomes during meiotic prophase.

## Supporting information

spotMAX parameter

## Acknowledgments

We thank members of the Köohler and Villeneuve lab, especially Celja Uebel, for critically reading the manuscript and help with data representation. We thank Luke Lavis (Janelia Research Campus) and Claire Deo (EMBL) for the JF639-HaloTag ligand, and Kurt Schmoller and Francesco Padovani (both Helmholtz Institute Munich) for help with spotMAX. We thank the Advanced Light Microscopy Facility (ALMF) at the European Molecular Biology Laboratory (EMBL) and Evident for support, IT and HPC resources at the EMBL in Heidelberg for providing essential computational infrastructure, the Protein Expression and Purification Core Facility at the EMBL for providing purified Proteinase K for this study, and the Media and Lab Kitchen at the EMBL in Heidelberg for providing media, solutions and sterilized material for this study. We also acknowledge the access and services provided by the Imaging Centre at the European Molecular Biology Laboratory (EMBL IC), generously supported by the Boehringer Ingelheim Foundation. Some strains were provided by the CGC, which is funded by NIH O”ce of Research Infrastructure Programs (P40 OD010440).

## Author contributions

I.C. performed all experiments and data analyses except for chromosome spread experiments, and wrote the original draft of the manuscript. A.W. carried out and analyzed the chromosome spread experiments and contributed to writing. Y.-L.W. assisted with SMLM data analysis and contributed to manuscript review and editing. E.B.D. and L.S. assisted with live imaging experiments and participated in review and editing. C.P.L. generated *C. elegans* strains and contributed to manuscript review and editing. F.H. and A.G. contributed to interference length analysis, with A.G. also assisting in review and editing. A.V., J.R., and S.K. supervised the research, acquired funding, and contributed to data analysis, writing of the original draft, and manuscript review and editing.

