## Supplement for "Multi-step implementation of meiotic crossover patterning"

### Supplementary Figures

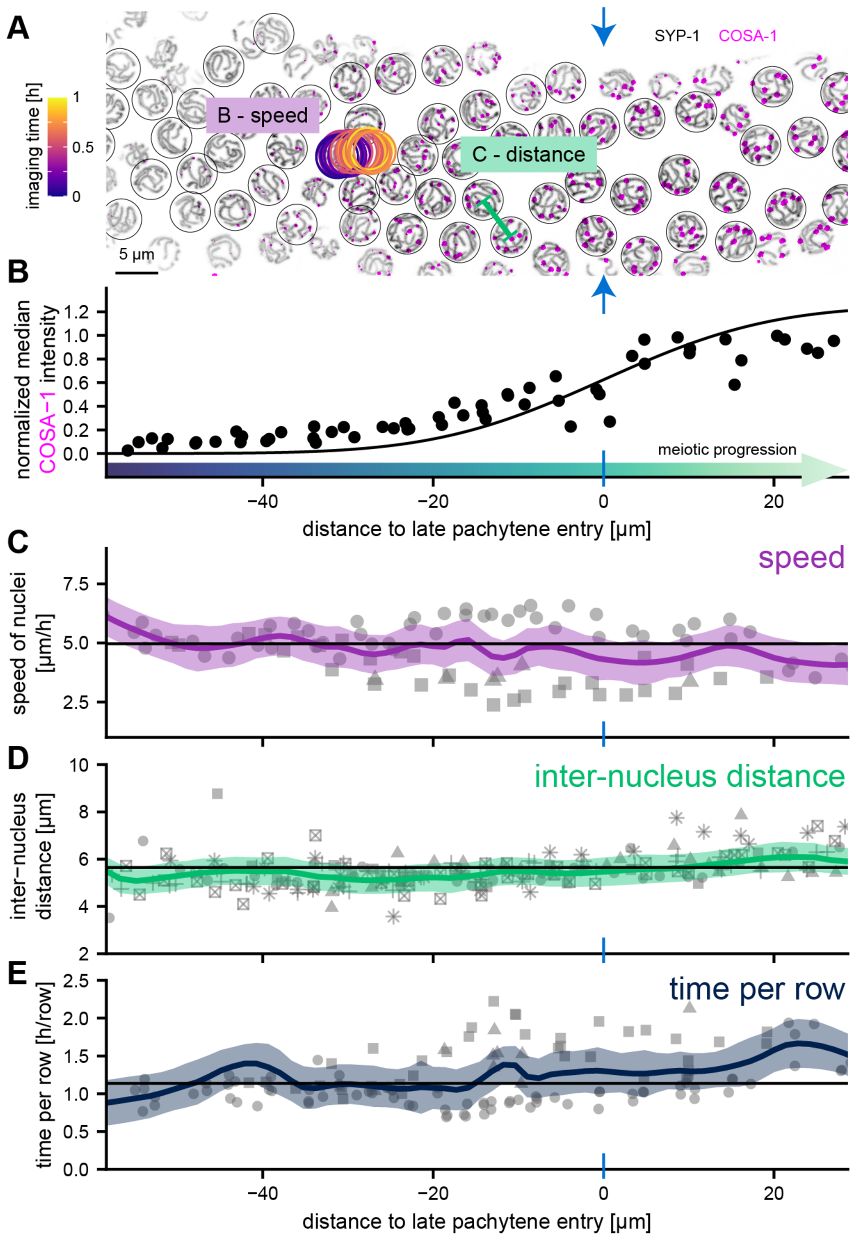


**Figure S1. Real-time imaging of meiotic progression in *C. elegans*.** (A) Maximum intensity projection of a snapshot (t = 0 min) showing meiotic nuclei labeled with SYP-1::mNeonGreen and Halo::COSA-1. Circles mark individual nuclei at their centroid positions. Colored circles highlight the position of a single nucleus over time relative to the point of entry into late pachytene (blue arrows). Row spacing is estimated from nearest-neighbor distances (example shown in green).

**Figure S1. Real-time imaging of meiotic progression in *C. elegans* (*continued*).** (B) Median intensities of COSA-1 foci for each nucleus were used to fit the point of entry into late pachytene (see Methods). (C) Nuclear speed was estimated from linear fits to trajectories of nuclei tracked for at least 70% of frames over $\geq$45 min (1 frame/min). (D) Nearest-neighbor distances as a proxy for row-to-row distances remain approximately constant throughout pachytene. (E) Time spent per row was calculated from nuclear speed and row-to-row distance. B-D: Different shapes represent individual animals; black lines indicate mean values; colored lines show loess fits with standard error.

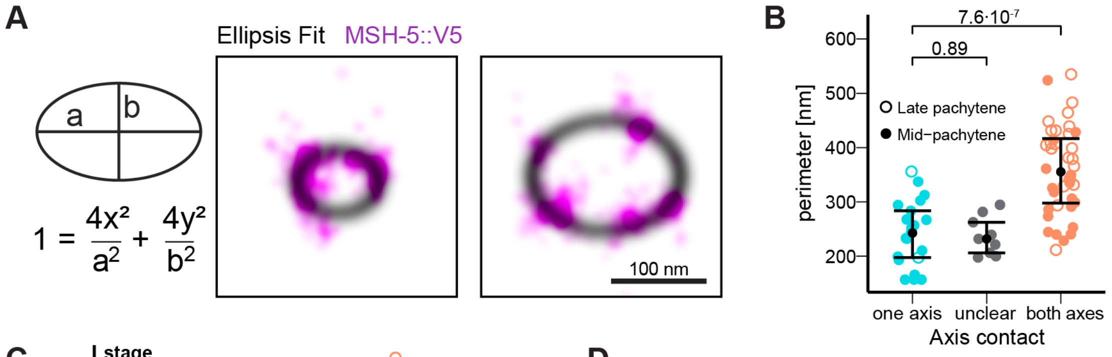


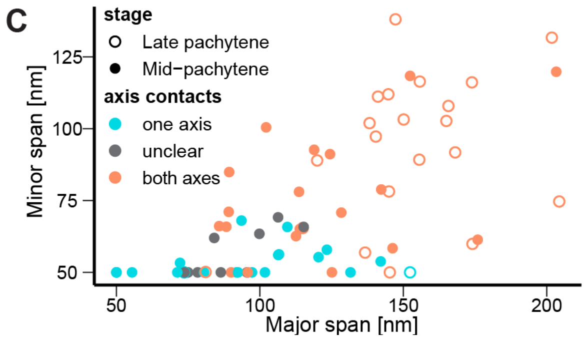


D

| Stage | #Total | Axis contacts | #Foci |
|---|---|---|---|
| Late pachytene | 23 | both axes | 21 |
| | | one axis | 2 |
| | | unclear | 0 |
| Mid-pachytene | 47 | both axes | 19 |
| | | one axis | 19 |
| | | unclear | 9 |

**Figure S2. Single-Molecule Localization Microscopy reveals distinct ultrastructures of MSH-5 recombination intermediates.** (A) Two examples of the ellipsis fit to the 3D-SMLM localizations of immunostained MSH-5::V5 that differ in the size of their respective ultrastructure. (B) Comparing the size (perimeter of the fitted ellipsis) of the ultrastructure of the individual MSH-5::V5 foci, depending on whether their respective localizations contact one or both chromosome axes. Color corresponds to the group as indicated on the *x-axis*. Error bars are median ± IQR. P-values are calculated by a Mann-Whitney test and adjusted using the Benjamini-Hochberg correction. (C) Distribution of the major and minor spans (described in A as "a" and "b") of the fitted ellipsis model across all analyzed foci (n=70). (D) Comparing the number of foci belonging to each class within late and mid-pachytene nuclei.

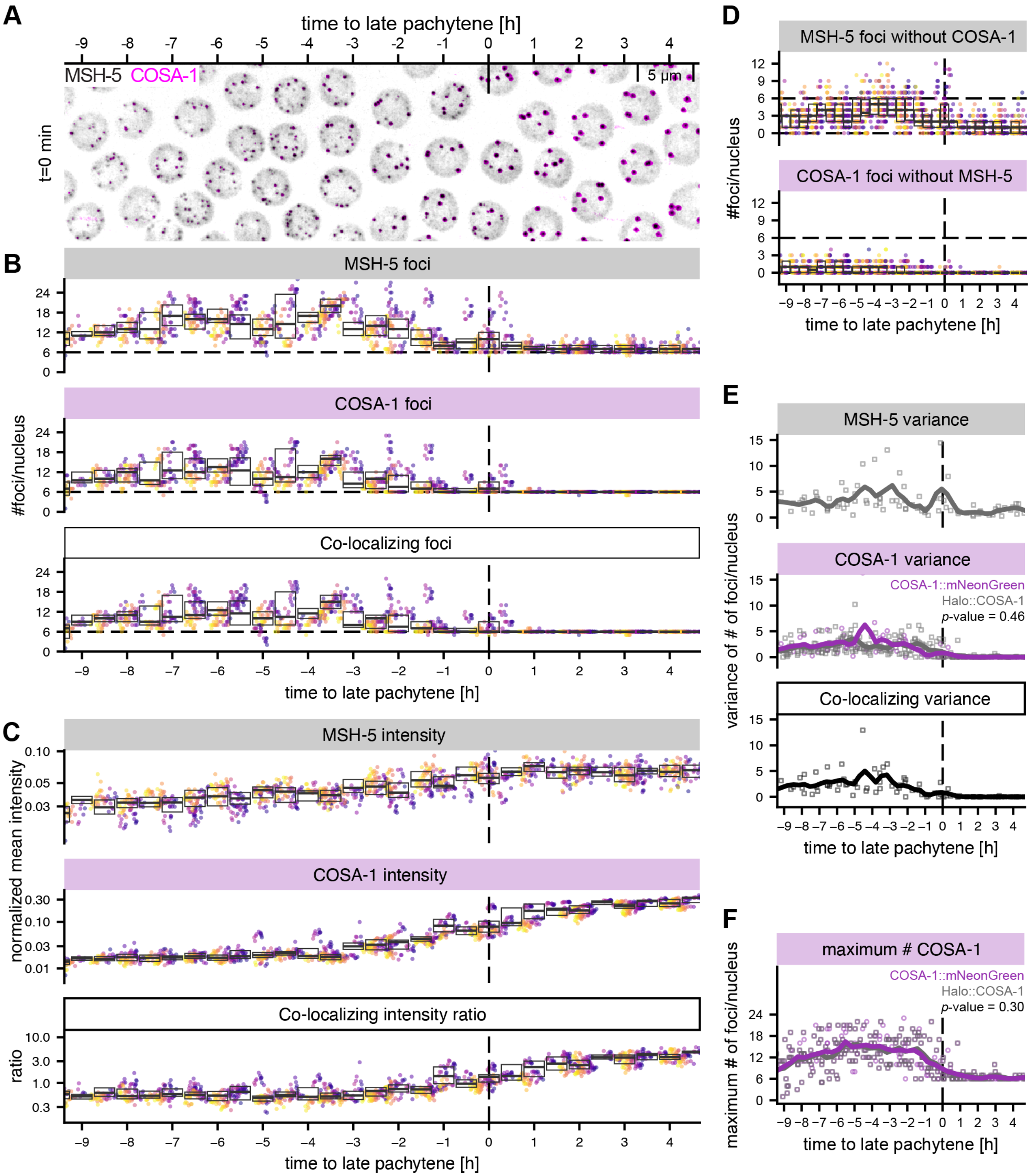


Figure S3. *In vivo* imaging of crossover designation.

**Figure S3. *In vivo* imaging of crossover designation (*continued*).** (A) Maximum intensity projection of Halo::MSH-5 (gray) and COSA-1::mNeonGreen at a single time point. (B) The number of MSH-5 (top), COSA-1 (center), and co-localizing (bottom) foci per nucleus first increases and then decreases to about six at the entry into late pachytene. Boxplots show interquartile ranges for nuclei at certain stages, single points show single nuclei with colors indicating imaging time (also applies to C and D). (C) The intensity of MSH-5 and COSA-1 increases during pachytene. The inflection point for COSA-1 intensity defines the point of entry into late pachytene for each time point (dashed lines in B-F). (D) The number of MSH-5 (top) and COSA-1 (bottom) foci that do not colocalize with the other marker are shown. Only a small number of COSA-1 foci do not overlap with MSH-5. 131 nuclei from 2 animals are analyzed in (B-D). (E) The variance of the number of foci per nucleus is high in mid-pachytene and low in late pachytene. Comparing the variance of the number of foci per nucleus and (F) the maximum number of foci shows no significant differences between COSA-1::mNeonGreen (131 nuclei from 2 gonads) and Halo::COSA-1 (336 nuclei from 6 gonads) behavior. p-values in E and F were calculated using generalized additive mixed models (see Methods).

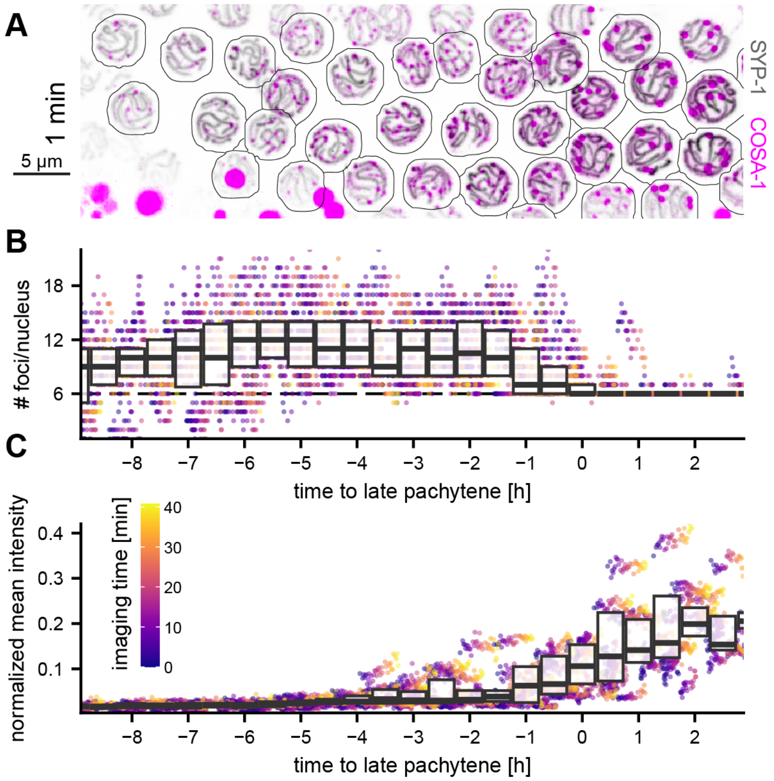


**Figure S4. COSA-1 focus intensity rises only after the number of foci per nucleus drops to six.** (A) Maximum intensity projection of SYP-1::mNeonGreen and Halo::COSA-1. Outlines of nuclei masks are shown as black lines. (B) Quantification of the number of COSA-1 foci. Boxplots show interquartile ranges for nuclei at specified positions imaged for 40 min, and points are measurements from individual cells and time points (scale bar for time in C). (C) The intensity of foci increases during pachytene. Boxplots show interquartile ranges of all measurements at specified positions and colored points show individual measurements with colors denoting imaging time. Data from 336 nuclei from 6 different animals are shown in B and C.

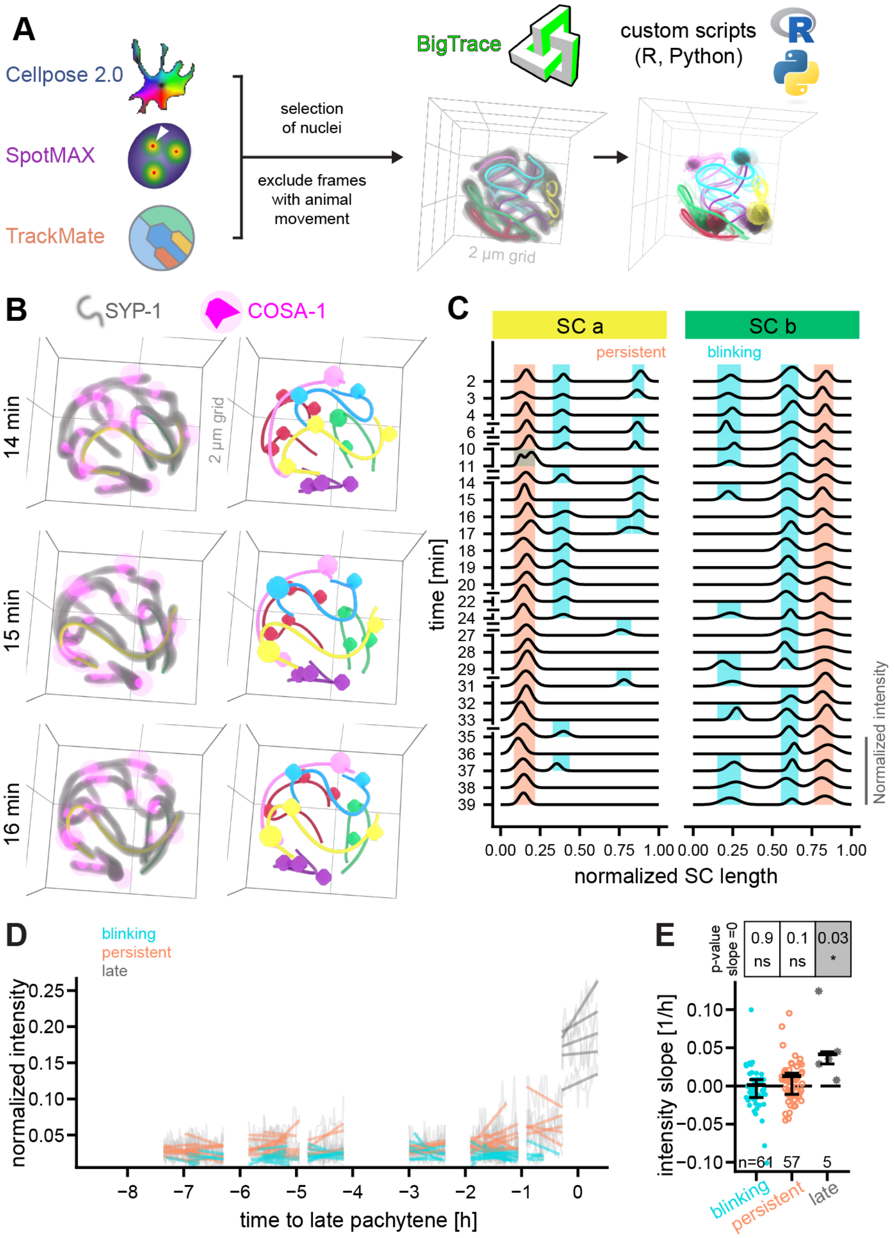


**Figure S5. Tracing of individual chromosomes and tracking the fate of recombination intermediates reveals two populations of COSA-1 foci.** (A) Image analysis pipeline to determine the dynamics of individual COSA-1 foci. After nucleus segmentation, spot detection, and tracking of all nuclei, a subset is selected for further analysis. Frames that show animal movement during the acquisitions are excluded from further analysis. Chromosomes are traced manually using BigTrace, and COSA-1 foci are mapped to the closest trace using custom scripts as described in the Methods. Visualizations are rendered using Blender and the Microscopy Nodes add-on.

**Figure S5. Tracing of individual chromosomes and tracking the fate of recombination intermediates reveals two populations of COSA-1 foci (*continued*).** (B) 3D renders of a single nucleus (left) and corresponding chromosome traces (right, colored lines) and associated spots (colored spheres) are shown. (C) Intensity profiles of COSA-1 foci along individual chromosomes (colors as in B) reveal two distinct types: persistent foci (orange), which are consistently present across all analyzable frames, and blinking foci (cyan), which transiently appear and disappear. (D) The intensity of individual foci only starts to increase in late pachytene. Light gray lines show fluctuations of individual foci and colored lines are linear fits (colored by type of foci). Slopes for individual foci are shown in (E). Intensities only increase significantly for late foci (one-sided Wilcoxon signed-rank test). The number of foci analyzed in D and E is given as n.

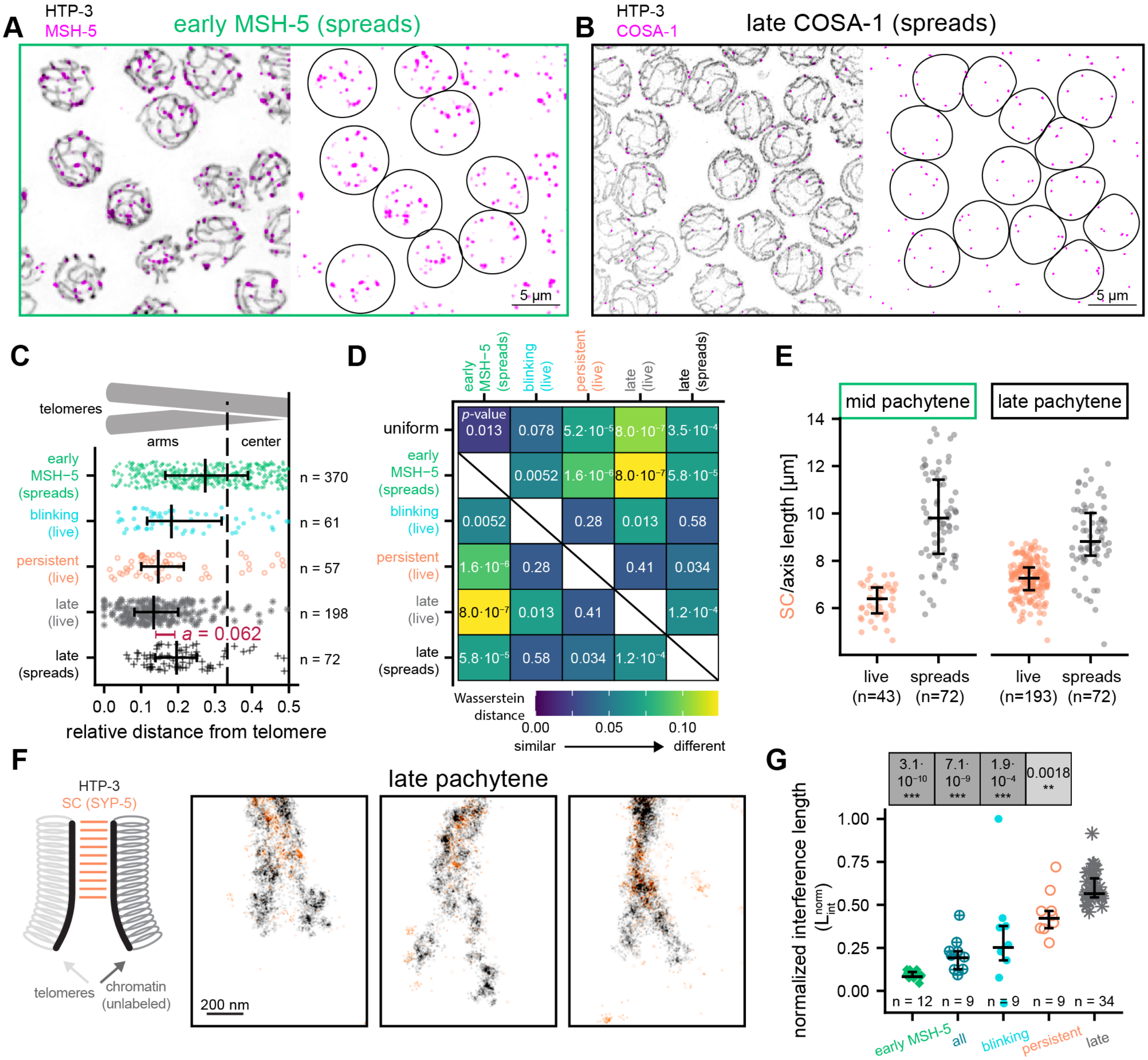


**Figure S6. Comparison and alignment of recombination focus positions between experimental approaches.** (A-B) Fluorescence images of chromosome spreads of mid-pachytene nuclei stained for chromosome axes (HTP-3) and early MSH-5 foci (A) or late pachytene COSA-1 foci (B) used for tracing positions of foci along axes.

**Figure S6. Comparison and alignment of recombination focus positions between experimental approaches (*continued*).**

(C) Distributions of foci along chromosomes, showing that persistent COSA-1 and late pachytene COSA-1 foci, observed in live imaging and spreads, are enriched on chromosome arms, while blinking foci and early MSH-5 foci in spreads show more random distributions. Due to methodological differences, the positions of late COSA-1 foci assessed in chromosome spreads were shifted relative to our measurements of late COSA-1 foci in live imaging. Consequently, the positions of foci from spreads $d$ were corrected for Fig. 4B using the transformation $d_{\text{corrected}} = 0.5 \times \frac{(d-a)}{(0.5-a)}$, where $a = 0.062$ (red) is the shift between the median positions of late COSA-1 foci in spreads and in live imaging data. The same correction was applied for positions of early MSH-5 foci in Fig. 4B. (D) Heat map showing pairwise Wasserstein distances, where color represents the degree of shift between the (uncorrected) distributions. Numbers are p-values for different comparisons from Kolmogorov-Smirnov tests after correction using the Benjamini-Hochberg method. (E) Length measurements for chromosomes are significantly longer in spread preparations than in live-imaging conditions. Mann–Whitney tests (corrected using the Benjamini–Hochberg method) show $p$-values $< 10^{-6}$ for all comparisons, except for mid- vs. late-pachytene spread chromosomes ($p = 0.009$). n indicates the number of traced chromosomes. (F) Super-resolution SMLM images of late pachytene chromosomes [90], showing that the SC central region does not extend all the way to the ends of the chromosome axes, leaving small regions near telomeres unsynapsed (also: [95]). This feature may contribute to the apparent shift in focus positions discussed above, as a chromosome axis marker was used for chromosome tracing in spread preparations, whereas SC central region markers were used for chromosome tracing in live imaging. These two features (E-F) and differences in tracing methods (see Methods) cause the shift between the two measurements as shown in (C). (G) Graphs show normalized interference lengths calculated using the method of [38]. Error bars show median $\pm$ interquartile ranges. Points show interference lengths for individual nuclei (n). $p$-values for comparisons to late COSA-1 foci were estimated by Mann-Whitney tests and corrected using the Benjamini-Hochberg method.

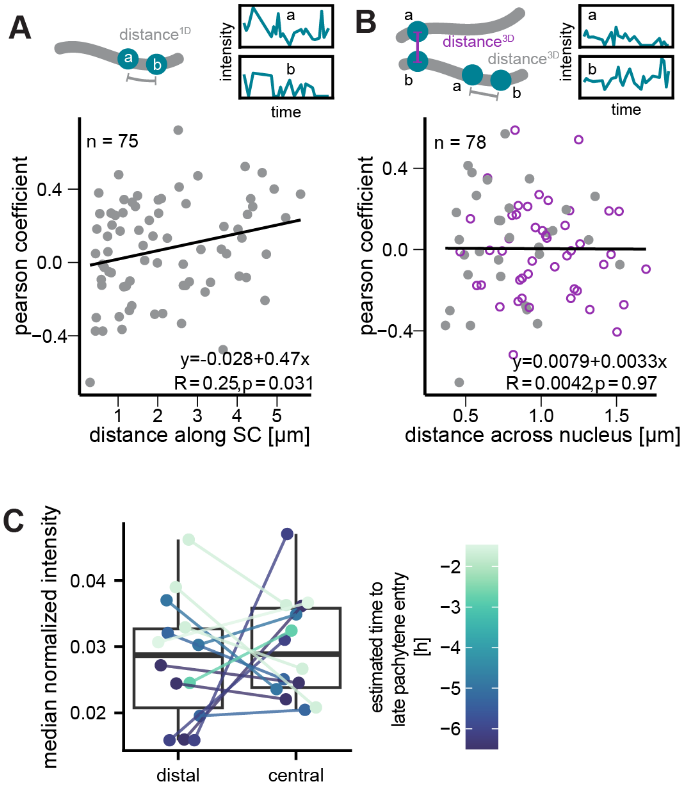


**Figure S7. Fluorescence intensities of neighboring foci do not correlate.** (A) The intensities of neighboring foci along individual SCs are not negatively correlated. Individual points show cross-correlations between intensities of neighboring foci located on the same SC at a distance$^{1D}$ as shown in the cartoon. (B) Similarly, the intensities of nearest-neighbor foci in the nucleus (distance$^{3D}$, cartoon) are not correlated. Gray spheres indicate that the nearest neighbor is on the same chromosome, purple open spheres show that the nearest neighbor is on a different chromosome. (C) For chromosomes containing two persistent foci, the median focus intensity of the more distal focus is not different from the more central focus (Mann-Whitney test p-value = 0.5; n=14 foci each). Colors of points show the position of the chromosomes in the germline; boxplots show interquartile ranges, and whiskers show extreme values.

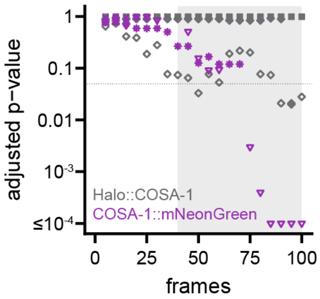


**Figure S8. COSA-1 foci exhibit minimal photobleaching effects during the first 40 frames.** Photobleaching can lead to a decline in the number of detected spots over time; however, the number of spots per nucleus remains largely stable for at least 40 frames across experiments (n=10). Therefore, analysis was restricted to the first 40 frames, although most experiments exhibited even less bleaching (gray area).

#### Supplemental tables

**Table S1.** *Caenorhabditis elegans* strains

| Strain ID | Strain | Source |
|---|---|---|
| **N2** | wild-type isolate, Bristol | CGC |
| **MT21793** | *gur-3(ok2245) lite-1(ce314)* X | CGC;<br>Bhatla and Horvitz,<br>2015 [66] |
| **SMN260** | *cosa-1(ske25[HaloTag::cosa-1])* III | Padovani et al., 2024 [33] |
| **SMN333** | *cosa-1(ske21[cosa-1::mNeonGreen])* III | Padovani et al., 2024 [33] |
| **SMN434** | *msh-5(ske50[msh-5::HaloTag9])* IV | This study |
| **SMN454** | *cosa-1(ske21[cosa-1::mNeonGreen])* III ; *msh-5(ske50[msh-5::HaloTag9])* IV | This study |
| **SMN256** | *cosa-1(ske25[HaloTag::cosa-1])* III; *syp-1(ske17[syp-1::mNeonGreen])* V;<br>*gur-3(ok2245) lite-1(ce314)* X | This study |
| **SMN311** | *htp-3(ske16-1[htp-3::HA])* I; *cosa-1(ske25[HaloTag::cosa-1])* III; *msh-5(ske49[msh-5::V5])* IV | This study |
| **AV842** | *meIs8*[unc-119(+); Ppie-1::gfp::cosa-1] II; *cosa-1(tm3298)* III; *him-6(jf93[him-6::HA]* IV | Jagut et al., 2016 [96] |
| **AV1445** | *opIs263*[rpa-1p::rpa-1::YFP + unc-119(+)]; *him-6(jf93[him-6::HA]* IV | Derived from<br>Stergiou et al., 2011 and<br>Jagut et al., 2016 [97, 96] |
| **SMN351** | *htp-3(ske16-1[htp-3::HA])* I; *cosa-1(ske25[HaloTag::cosa-1])* III; *rad-51(ske48[rad-51::V5])* IV | This study |
| **SMN348** | *htp-3(ske16-1[htp-3::HA]) zhp-3(ske71[zhp-3::V5])* I; *cosa-1(ske25[HaloTag::cosa-1])* III | This study |

**Table S2.** Repair templates for CRISPR/Cas9-mediated genome editing

| Allele (genotype) | Repair template sequence | Description |
|---|---|---|
| ***ske17* (*syp-1::mNeonGreen*)** | 5’-cgtgcggccgatcgtcttaaaacgagagccgaagctcatactgcagat<br>gtaagaagaaaacgcggagggaagaaagtctccaagggagaggaggacaa<br>catggcctcccttccagccacccacgagcttcacatcttcggatccatca<br>acggagtcgacttcgacatggtcggacagggaaccggaaacccaaacgac<br>ggatacgaggagcttaaccttaagtccaccaaggtaagtttatctaaaaa<br>gtttttcattcaaaatgtgtaaaaattcatttaaaataacccaaaaatca<br>ttaatcctcgatattttcagggagaccttcaattctccccatggatcctt<br>gtcccacacatcggatacggattccaccagtaccttccatacccagacgg<br>aatgtccccattccaagccgccatggtcgacggatccggataccaagtcc<br>accgcaccatgcaattcgaggacggagcctcccttaccgtcaactaccgc<br>tacacctacgagggatcccacatcaagggagaggcccaagtcaagggaac<br>cggattcccagccgacggaccagtcatgaccaactcccttaccgccgccg<br>actggtgccgctccaagaagacctacccaaacgacaagaccatcatctcc<br>accttcaagtggtcctacaccaccggaaacggaaagcgctaccgctccac<br>cgcccgcaccacctacaccttcgccaagccaatggccgccaactacctta<br>agaaccagccaatgtacgtcttccgcaagaccgagcttaagcactccaag<br>accgagcttaacttcaaggagtggcagaaggccttcaccgacgtcatggg<br>aatggacgagctttacaagggaggtggctaatgtgtgtgtggggaagaaa<br>cgactatgtaccatttcaatcttgtgctatttttttttgtgtttttagag<br>ttt-3’ | Initial gBlock ordered from IDT was cloned into pMiniT 2.0 (NEB® PCR Cloning Kit, New England Biolabs, Inc., E1202S). The single stranded template (final concentration **180 ng/µl** in the injection mix) was generated by PCR using high-fidelity DNA polymerase (Q5® High-Fidelity DNA Polymerase, New England Biolabs, Inc., M0491S) and the following primers: F: 5’-cgtgcggccgatcgtcttaaa acgagagccgaagctcatactgcaga tgtaagaagaaaacgcggagggaag aaagtctccaagggagaggagga-3’; R: 5’-aaactctaaaaacacaaaaaa aaatagcacaagattgaaatggtaca tagtcgtttcttccccacacacaca ttagccacctcccttgtaaagct-3’; single stranded DNA was generated by a primer extension PCR using the biotinylated primer: 5’-(btn)cgtgcgg ccgatcgtctt-3’ |

Table S2 – *Continued on the next page*

Table S2 – *Continued from the previous page*

| **Allele (genotype)** | **Repair template sequence** | **Description** |
|---|---|---|
| ***ske49* (*msh-5::V5*)** | 5’-ggaagatgatgagttcttgaagagtttccttgaaacagaaggatctct acatattgacacaagcgccgatgagacgatagatcgttcgaaaagaagtg gaaagccaatcccaaacccacttcttggactcgactccacctaattttat ataattagaatttcgtatttcttgtaatgttcaatcttgttcaatgtatt tcgttttcgtttttaaattatttatagcgcattttaataaatactatttt ttaaaactcgtgcaatagttcacaacgaagtttaactgcgatttttttaca aaaaatacagatttttttggttttcaaaagtagctgatctgcaattattgc actttctgaaagtatcgaattaatgaatgcaacaagaagttccagacgcg cacttttgcaattcttgcgcgaaatatactgtacccgggc-3’ | Initial gBlock ordered from IDT was cloned into pMiniT 2.0 (NEB® PCR Cloning Kit, New England Biolabs, Inc. E1202S). Injected template (final concentration **62 ng/µl**) was generated by PCR using high-fidelity DNA polymerase (Q5® High-Fidelity DNA Polymerase, New England Biolabs, Inc., M0491S) and following primers: F: 5’-ggaagatgatgagttcttgaaga-3’; R: 5’-gcccgggtacagtatatttc-3’ |

Table S2 – *Continued on the next page*

Table S2 – *Continued from the previous page*

| Allele (genotype) | Repair template sequence | Description |
|---|---|---|
| ***ske50* (*msh-5::HaloTag9*)** | 5'-ggaagatgatgagttcttgaagagtttccttgaaacagaaggatctct acatattgacacaagcgccgatgagacgatagatcgttcgaaaagaagtg gagctggaatcggaaccggattcccattcgacccacactacgtcgaggtc cttggagagcgcatgcactacgtcgacgtcggaccacgcgacggaacccc agtccttttccttcacggaaacccaacctcctcctacgtctggcgcaaca tcatcccacacgtcgccccaacccaccgctgcatcgccccagaccttatc ggaatgggaaagtccgacaagccagaccttggatacttcttcgacgacca cgtccgtttcatggacgccttcatcgaggcccttggacttgaggaggtcg tccttgtcatccacgactggggatccgcccttggattccactgggccaag cgcaacccagagcgcgtcaaggtaagtttaaacatatatatactaactaa ccctgattatttaaattttcagggaatcgccttcatggagttcatccgcc caatcccaacctgggacgagtggccagagttcgcccgcgagaccttccaa gccttccgcaccaccgacgtcggacgtaagcttatcatcgaccacaacgt cttcatcgagggaacccttcgtatgggagtcgtccgtccacttaccgagg tcgagatggaccactaccgcgagccattccttaacccagtcgaccgcgag ccactttggcgcttcccaaacgagcttccaatcgccggagagccagccaa catcgtcgcccttgtcgaggagtacatggactggcttcaccaatccccag tcccaaagcttcttttctggggaaccccaggagtccttatcccaccagcc gaggccgcccgtcttgccaagtcccttccaaactgcaaggtaagtttaaa cagttcggtactaactaaccatacatatttaaattttcaggccgtcgaca tcggaccaggacttaaccttcttcaagaggacaacccagaccttatcgga tccgagatcgcccgttggctttccacccttgagatctaattttatataat tagaatttcgtatttcttgtaatgttcaatcttgttcaatgtatttcgtt ttcgtttttaaattatttatagcgcattttaataaatactatttttttaaa actcgtgcaatagttcacaacgaagtttaactgcgatttttttacaaaaaa tacagatttttttggttttcaaaagtagctgatctgcaattattgcacttt ctgaaagtatcgaattaatgaatgcaacaagaagttccagacgcgcactt ttgcaattcttgcgcgaaatatactgtacccgggc-3' | Initial gBlock ordered from IDT was cloned into pMiniT 2.0 (NEB® PCR Cloning Kit, New England Biolabs, Inc., E1202S). Injected template (final concentration **40 ng/µl**) was generated by PCR using high-fidelity DNA polymerase (Q5® High-Fidelity DNA Polymerase, New England Biolabs, Inc., M0491S) and following primers: F: 5'-ggaagatgatgagttcttgaaga-3'; R: 5'-gcccgggtacagtatatttc-3'; Before it was added into the injection mixture template was melted according to protocol described in Ghanta and Mello, 2021 [58]. |

**Table S3.** Sequences of CRISPR-RNA (crRNA), genotyping information, and sequencing primers for each CRISPR/Cas9-mediated genome editing

| Allele (genotype) | crRNA | Genotyping primers and fragment sizes | Sequencing primers |
|---|---|---|---|
| ***ske17*** **(*syp-1::mNeonGreen*)** | 5'-gatgttcgccgaaagagagg-3' | F:5'-cgatatcgtggaatccgact-3';<br>R:5'-ccaatttgtcggggagtttt-3';<br>WT, 198 bp; Edit, 1210 bp | F:5'-cgatatcgtggaatccgact-3';<br>R:5'-ccaatttgtcggggagtttt-3'; |
| ***ske49*** **(*msh-5::V5*)** | Two crRNAs were used simultaneously:<br>5'-catcggcgctcgtatcgata-3' and 5'-tgcgcgtctggaacttcttg-3' | F:5'-gtcaaacgacgatgaggaag-3';<br>R:5'-cgttgtgaactattgcacgag-3';<br>WT, 248 bp; Edit, 290 bp | F:5'-gttcgatttccccgagca-3';<br>R:5'-taccttcacaactcttccatcg-3'; |
| ***ske50*** **(*msh-5::HaloTag9*)** | Two crRNAs were used simultaneously:<br>5'-catcggcgctcgtatcgata-3';<br>5'-tgcgcgtctggaacttcttg-3' | **Set1:**<br>F:5'-gttcgatttccccgagca-3';<br>R:5'-gggcgatgcagcgg-3';<br>WT, no amplification; Edit, 474 bp;<br>**Set2:**<br>F:5'-gttcgatttccccgagca-3';<br>R:5'-cgttgtgaactattgcacgag-3';<br>WT, 420 bp; Edit, 1407 bp | F:5'-gttcgatttccccgagca-3';<br>R:5'-taccttcacaactcttccatcg-3' |

**Table S4.** Brood counts for the *Caenorhabditis elegans* strains

| **Strain ID (genotype)** | **# eggs** | **# males** | **# adults** | **Viability of progeny [%]** | ***p*-value[a]** | **Incidence of males [%]** | ***p*-value[a]** |
|---|---|---|---|---|---|---|---|
| **N2** (wild-type isolate, Bristol) | 3221 (11) | 4 | 4009 | 121.6 | 0.73 (ns) | 0.1 | 0.009 (**) |
| **SMN260** (*cosa-1*(*ske25*[*HaloTag::cosa-1*]) III) | 1548 (6) | 8 | 1612 | 103.5 | *reference group* | 0.46 | *reference group* |
| **SMN434** (*msh-5*(*ske50*[*msh-5::HaloTag9*]) IV) | 1583 (6) | 5 | 1630 | 102.9 | 1.00 (ns) | 0.30 | 0.23 (ns) |
| **SMN454** (*cosa-1*(*ske21*[*cosa-1::mNeonGreen*]) III; *msh-5*(*ske50*[*msh-5::HaloTag9*]) IV) | 1680 (6) | 6 | 1710 | 101.9 | 0.87 (ns) | 0.38 | 0.38 (ns) |
| **SMN256** (*cosa-1*(*ske25*[*HaloTag::cosa-1*]) III; *syp-1*(*ske17*[*syp-1::mNeonGreen*]) V; *gur-3*(*ok2245*) X; *lite-1*(*ce314*) X) | 1705 (6) | 7 | 1711 | 100.6 | 0.59 (ns) | 0.41 | 0.75 (ns) |
| **SMN311** (*htp-3*(*ske16-1*[*htp-3::ha*]) I; *cosa-1*(*ske25*[*HaloTag::cosa-1*]) III; *msh-5*(*ske49*[*msh-5::V5*]) IV) | 1404 (5) | 7 | 1511 | 107.8 | 0.31 (ns) | 0.46 | 0.75 (ns) |

[a]Wilcoxon rank sum test using the *ggpubr* R package (version 0.6.0). Statistical significance: ns ($p > 0.05$), * ($p <= 0.05$), ** ($p <= 0.01$).

**Table S5. Typical TrackMate parameters used to track meiotic nuclei during timelapse movies of *Caenorhabditis elegans* germline**

| TrackMate parameter | Value |
|---|---|
| ALLOW_TRACK_SPLITTING | FALSE |
| ALLOW_TRACK_MERGING | FALSE |
| LINKING_MAX_DISTANCE [µm][b] | 7-10 |
| GAP_CLOSING_MAX_DISTANCE [µm][b] | 10-20 |
| MAX_FRAME_GAP[b] | 3-5 |
| LINKING_FEATURE_PENALTIES | {'POSITION_Y': 2.,'POSITION_Z': 15.,'RADIUS': 7.} |
| GAP_CLOSING_FEATURE_PENALTIES | {'POSITION_Y': 2., 'POSITION_Z': 7.,'RADIUS': 5.} |

[b] A range of typically used values is listed.

**Table S6.** Immunostaining details for imaging of the germline spreads

| Stage | Strain | DeltaVision OMX™ Blaze module | Primary antibody | Secondary antibody | Target |
|---|---|---|---|---|---|
| **Mid-pachytene** | **AV1445** | **Widefield imaging with deconvolution** | Chicken polyclonal anti-HTP-3 (MacQueen et al., 2005 [98]) | Goat polyclonal anti-chicken AlexaFluor 405 conjugated (Abcam, ab175675) | HTP-3[c] |
| | | | Alpaca monoclonal anti GFP Atto488 conjugated (Chromotek, gba488-100) | N/A | RPA-1::YFP |
| | | | Mouse monoclonal anti-HA (Biolegend, 901501) | Goat polyclonal anti-mouse AlexaFluor 555 conjugated (Molecular Probes, A-21422) | HIM-6::HA |
| | | | Rabbit anti-MSH-5Pi (Haversat et al., 2022 [99]) | Goat polyclonal anti-mouse AlexaFluor 647 conjugated (Molecular Probes, A-21235) | MSH-5[d] |
| **Late Pachytene** | **AV842** | **3D structured illumination microscopy** | Alpaca monoclonal anti GFP Atto488 conjugated (Chromotek, gba488-100) | N/A | GFP::COSA-1[d] |
| | | | Guinea pig polyclonal anti-SYP-1 (MacQueen et al., 2002 [100]) | Goat polyclonal anti-guinea pig Alexa 555 conjugated (Thermo Fisher Scientific, A21435) | SYP-1 |
| | | | Chicken polyclonal anti-HTP-3 (MacQueen et al., 2005 [98]) | Goat polyclonal anti-chicken AlexaFluor 647 conjugated (Thermo Fisher Scientific, A21449) | HTP-3[c] |

[c] Signal used to trace the chromosomes.
[d] Signal used to identify the foci.

**Table S7. LocMoFit parameters used to initialize ellipsis fit**

| name | initial value | fix | lb | ub | type | min | max | unit |
|---|---|---|---|---|---|---|---|---|
| x | 0 | 0 | -50 | 50 | lPar | -200 | 200 | nm |
| y | 0 | 0 | -50 | 50 | lPar | -200 | 200 | nm |
| zrot | 0 | 0 | -Inf | Inf | lPar | -180 | 180 | ° |
| variation | 8 | 1 | 0 | 10 | lPar | 0 | 20 | nm |
| xscale | 1 | 1 | 1 | 1 | lPar | 1 | 1 | |
| yscale | 1 | 1 | 1 | 1 | lPar | 1 | 1 | |
| z | 0 | 0 | -20 | 20 | lPar | -400 | 400 | nm |
| xrot | 0 | 0 | -Inf | Inf | lPar | -180 | 180 | ° |
| yrot | 0 | 0 | -Inf | Inf | lPar | -180 | 180 | ° |
| zscale | 1.5 | 1 | 1 | 1 | lPar | 1 | 1 | |
| semiaxis a | 50 | 0 | -Inf | Inf | mPar | 25 | 150 | nm |
| semiaxis b | 50 | 0 | -Inf | Inf | mPar | 25 | 150 | nm |

#### Captions for videos

**Video S1. Ultrastructures of MSH-5 recombination nodules in late pachytene.** 3D rendering of images shown in Fig. 1D, late pachytene.

**Video S2. Ultrastructures of MSH-5 recombination nodules show crossover-like features in mid-pachytene.** 3D rendering of images shown in Fig. 1D, mid-pachytene.

**Video S3. Time-lapse images reveal dynamic association of MSH-5 and COSA-1 with recombination nodules.** Time-lapse movie of Fig. 2B.

**Video S4. High-resolution 3D renderings capture the dynamics of MSH-5 and COSA-1 in individual nuclei.** Movie corresponding to Fig. 2C. Shaded areas indicate spot fits obtained using spotMAX. Time frames with worm movement during image acquisition were excluded and thus do not display spotMAX results.

**Video S5. Time-lapse images of COSA-1 recombination nodules along individual chromosomes.** Timelapse Movie of Fig. 3A.

**Video S6. 3D renders of an individual nucleus show blinking and persistent COSA-1 foci in mid-pachytene.** Timelapse Movie of Fig. 3B.
