## Supplementary material for "Multi-step implementation of meiotic crossover patterning": spotMAX parameter: S1_COSA-1-mNG.ini

```
[File paths and channels]
Experiment folder path(s) to analyse = /g/koehler/ivana/Microsopy/ConfocalMicroscopy/SpinSR/240316_on_COSA-1-mNG_MSH-5-HT9-JF639/03_0_488i15_640i45_x200ms_TI60/TIFFs/Position_2/
Spots channel end name or path = 2
Cells segmentation end name or path = segm.npz
Reference channel end name or path = 
Spots channel segmentation end name or path = 
Ref. channel segmentation end name or path = 
Spots coordinates table end name or path = 
Table with lineage info end name or path = 
Run number = 10
Text to append at the end of the output files = 
File extension of the output tables = .h5

[METADATA]
Number of frames (SizeT) = 1
Analyse until frame number = 1
Number of z-slices (SizeZ) = 46
Pixel width (μm) = 0.065
Pixel height (μm) = 0.065
Voxel depth (μm) = 0.1999999999998181
Numerical aperture = 1.35
Spots reporter emission wavelength (nm) = 575.0
Spot minimum z-size (μm) = 0.51
Resolution multiplier in y- and x- direction = 1
Spot (z, y, x) minimum dimensions (radius) = 

[Pre-processing]
Aggregate cells prior analysis = False
Threshold only inside segmented objects = True
Remove hot pixels = True
Initial gaussian filter sigma = 1
Sharpen spots signal prior detection = True
Extend 3D input segm. objects in Z = (0, 0)

[Reference channel]
Segment reference channel = False
Keep only spots that are inside ref. channel mask = False
Use the ref. channel mask to determine background = False
Ref. channel is single object (e.g., nucleus) = False
Keep external touching objects intact = False
Ref. channel gaussian filter sigma = 0.75
Sigmas used to enhance network-like structures = 0.0
Ref. channel segmentation method = Thresholding
Ref. channel threshold function = threshold_otsu
Features for filtering ref. channel objects = 
Save reference channel features = False
Save reference channel segmentation masks = False
Save pre-processed reference channel image = False
Calculate reference channel network length = False

[Spots channel]
Spots segmentation method = Thresholding
Minimum size of spot segmentation mask = 8
Spot detection threshold function = threshold_triangle
Spots detection method = peak_local_max
Features and thresholds for filtering true spots = spot_vs_backgr_effect_size_hedge, 1.1
Local background ring width = 5 pixel
Optimise detection for high spot density = False
Compute spots size (fit gaussian peak(s)) = True
After spotFIT, drop spots that are too close = True
Merge spots pairs where single peak fits better = False
Maximum number of spot pairs to check = 11
Save spots segmentation masks = False
Save pre-processed spots image = False
Skip objects where segmentation failed = False

[SpotFIT]
Bounds interval for the x and y peak center coord. = 0.1
Bounds interval for the z peak center coord. = 0.2
Bounds for sigma in x-direction = 0.6*x_resolution_pxl/2.35, 1.4*x_resolution_pxl/2.35
Bounds for sigma in y-direction = 0.6*x_resolution_pxl/2.35, 1.4*x_resolution_pxl/2.35
Bounds for sigma in z-direction = 0.6*z_resolution_pxl/2.35, 1.4*z_resolution_pxl/2.35
Bounds for the peak amplitude = 0.0, spotsize_A_max
Bounds for the peak background level = spotsize_backgr_median - 0.01*spotsize_backgr_median, spotsize_backgr_median + 0.01*spotsize_backgr_median
Initial guess for sigma in x-direction = x_resolution_pxl/2.35
Initial guess for sigma in y-direction = x_resolution_pxl/2.35
Initial guess for sigma in z-direction = z_resolution_pxl/2.35
Initial guess for the peak amplitude = spotsize_A_max
Initial guess for the peak background level = spotsize_backgr_median

[Custom combined measurements]
Column name = 

[Configuration]
Folder path of the log file = /g/koehler/ivana/Microsopy/ConfocalMicroscopy/SpinSR/240316_on_COSA-1-mNG_MSH-5-HT9-JF639/03_0_488i15_640i45_x200ms_TI60/TIFFs/Position_2/_logs
Folder path of the final report = /g/koehler/ivana/Microsopy/ConfocalMicroscopy/SpinSR/240316_on_COSA-1-mNG_MSH-5-HT9-JF639/03_0_488i15_640i45_x200ms_TI60/TIFFs/Position_2//_logs
Filename of final report = 
Disable saving of the final report = True
Use default values for missing parameters = True
Stop analysis on critical error = False
Use CUDA-compatible GPU = False
Number of threads used by numba = 1
Reduce logging verbosity = True
```
